# Pericytes orchestrate a tumor-restraining microenvironment in glioblastoma

**DOI:** 10.1101/2024.08.26.609765

**Authors:** Sebastian Braun, Paulina Bolivar, Clara Oudenaarden, Jonas Sjölund, Matteo Bocci, Katja Harbst, Mehrnaz Safaee Talkhoncheh, Bengt Phung, Eugenia Cordero, Rebecca Rosberg, Elinn Johansson, Göran B Jönsson, Alexander Pietras, Kristian Pietras

**Author notes:** These authors contributed equally (Sebastian Braun, Paulina Bolivar).

## Abstract

Glioblastoma (GBM) is characterized by fast progression, an infiltrative growth pattern, and a high rate of relapse. A defining feature of GBM is the existence of spatially and functionally distinct cellular niches, i.e. a hypoxic niche, a leading-edge niche, and a perivascular niche, in which malignant cells engage in paracrine crosstalk with cell types comprising the tumor microenvironment. Here, by analysis of single-cell transcriptomic data of human GBM and transgenic mouse models of GBM, we unexpectedly identified pericytes, mural cells intimately associated with the endothelium, as the most active paracrine signaling hub within the tumor parenchyma. Exclusive signaling axes emanating from pericytes were received by endothelial cells, malignant cells, astrocytes, and immune cells. Depletion of pericytes through genetic engineering in several different transgenic and orthotopic mouse models of GBM demonstrated accelerated tumor progression, a disrupted blood-brain-barrier, and premature death of pericyte-poor mice. Mechanistic studies revealed that pericyte deficiency altered the cellular composition of GBM, remodeled the endothelium, and impacted on the immune cell landscape, exacerbating tumor cell invasion and immune suppression. Specifically, endothelial cells deprived of pericyte association altered their signaling programs, which in turn attracted perivascular, tumor-associated macrophages polarized towards an immune-suppressive phenotype. The recruited macrophages expressed Hepatocyte Growth Factor (HGF), which reinforced activation of its receptor tyrosine kinase MET on GBM cells harboring an extreme mesenchymal subtype driven by the key phenotypic regulator Fosl1 within hypoxic regions. Indeed, orthotopic implantation of isolated, MET-expressing GBM cells corroborated their superior tumor-initiating capability and invasive phenotype. In patients, low expression of a pericyte core gene signature was reduced in recurrent GBM, compared to primary tumors. Consistently, gene signatures for transcriptional programs of Fosl1^+^Met^+^ GBM cells were indicative of poor survival in human tumors, and spatial transcriptomics corroborated their superior invasive capacity. Taken together, we infer that the pericyte represents a critical modulator of GBM development by orchestrating a tumor-suppressive microenvironment; our findings thus highlight the importance of pericyte preservation in the face of current and future GBM therapies.

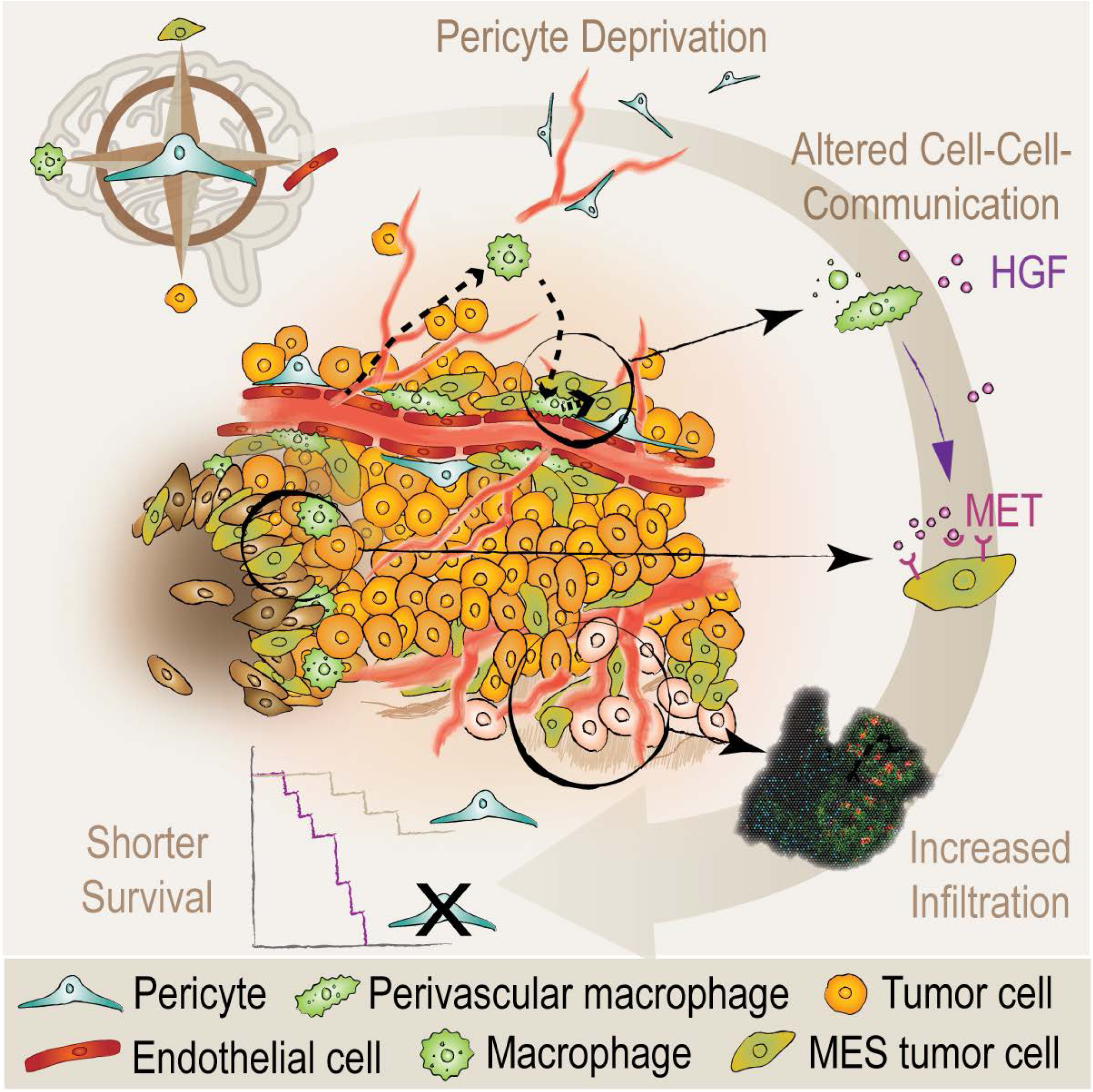

## Introduction

Glioblastoma (GBM) is an aggressive high-grade brain tumor, characterized by fast progression, an infiltrative growth pattern, and a high rate of relapse. Despite significant efforts over the last decades, GBM remains resistant to current post-surgical therapeutic approaches, such as chemotherapy with temozolomide, radiotherapy, or tumor-treating fields (Schaff and Mellinghoff, 2023; Stupp et al., 2005). The dismal prognosis of GBM is likely a result of diverse intrinsic features, including intra-tumoral heterogeneity, phenotypic plasticity, and a complex interplay between tumor cells and their microenvironment (Varn et al., 2022; Wang et al., 2022). The latter is manifested as a complex tumor ecosystem, consisting of interconnected but spatially and functionally distinct niches, such as the leading-edge niche, the hypoxic niche, and the perivascular niche (PVN) (Hambardzumyan and Bergers, 2015). Recently, it has become clear that glioma cells co-evolve with such niches and form specific molecular cell states that are adapted to the respective microenvironment (De Silva et al., 2023). Remarkably, due to a high degree of glioma cell intrinsic plasticity, gliomas can also react to and escape from environmental pressures such as lack of oxygen or therapeutically induced stress (Prager et al., 2020). However, on a higher level of granularity, our understanding of tumor, stromal and immune cell interactions is still inadequate, and the nature and sequence of the processes entailing glioma cell adaptation remain elusive.

The GBM vasculature is characterized by microvascular proliferation and functional abnormality (Diaz-Flores et al., 2021). It represents a crucial factor for tumor development and dissemination: glioma cells are presumably guided by chemotactic factors secreted by the vasculature, and contact-dependent interactions; processes recapitulating the physiologic migration patterns of glial progenitor cells that allow for fast and diffuse permeation of the surrounding brain parenchyma along blood vessels (Farin et al., 2006; Seano and Jain, 2020). Moreover, tumor-driving, stem-like glioma cells have been found to prosper along micro-vessels (Calabrese et al., 2007; Charles et al., 2011). Given the character of the PVN as a crosspoint of tumor and stromal cells, and a crucial infrastructure for brain infiltration, deepened insights on how the aberrant GBM vasculature contributes to glioma progression are much needed and may support the design of more effective therapies.

Pericytes, mesenchymal smooth muscle-like cells, own a key position in the PVN due to their location: they are embedded in the basal lamina of micro-vessels and in contact with both endothelial and tumor cells (Armulik et al., 2011). Whereas pericytes have been extensively described to exert functions as regulators of vascular morphogenesis, angiogenic quiescence, and blood-brain barrier patency, consequently maintaining vessel homeostasis in the physiologic brain context (Armulik et al., 2010; Bell et al., 2010), their role in brain tumors is still debated; pericytes have been linked to the induction of tumor immune-tolerance, shielding glioma cells from effective T-cell responses (Valdor et al., 2019), and have been suggested as drug targets to improve treatment of brain tumors (Zhou et al., 2017). Emerging evidence, however, demonstrates heterogeneity among the group of mural cells, and that pericytes are not the only mesenchymal stromal cells to reside at glioma capillaries (Andaloussi Mae et al., 2020; Jain et al., 2023; Oudenaarden et al., 2022), demanding a deeper characterization of glioma developmental processes related to true pericytes.

Here, by analysis of single-cell transcriptomic data of human GBM, we unexpectedly identified pericytes as the most active paracrine signaling hub within the tumor parenchyma. This finding prompted us to study the effects of pericyte depletion in orthotopic and transgenic mouse models of high-grade glioma using single-cell and spatial transcriptomics, as well as multiplex immunohistochemistry and mechanistic studies. We present evidence that pericytes mitigate glioma development by orchestrating a tumor-suppressive microenvironment, challenging the concept of pericytes as merely pro-oncogenic cells, and underscoring the importance of pericyte preservation in future GBM therapies.

## Results

### Pericytes represent a signaling hub in human GBM

We commenced our investigations with the goal of characterizing the heterogeneity of glioma, stromal and immune cells, and to better understand how the cells forming the tumor microenvironment (TME) communicate and interact within different niches. Therefore, we leveraged a public dataset, comprising single cell RNA sequencing (scRNA-seq) of 201,986 tumor, immune, and stromal cells, derived from two low-grade and 16 high-grade human glioma samples (Abdelfattah et al., 2022). In total, seven main cell clusters were annotated, based on well-known cell type markers, representing glioma cells, B-and T-cells, myeloid cells, endothelial cells, and oligodendrocytes (Figures 1A and S1A). A perivascular cell/pericyte cluster was defined based on the expression of established marker genes such as *S1PR3*, *ABCC9*, *CD248*, *RGS5*, *ANPEP*, *CSPG4*, *PDGFRβ*, *HIGD1B*, *NDUFA4L2*, *VTN* and *KCNJ8* (van Splunder et al., 2024) (Figure 1B). To learn more about the interactions between the different tumor, immune and stromal cell compartments, we applied two algorithms, CellChat and CellPhoneDB, that use different computational approaches to quantitatively characterize intercellular communication networks (Jin et al., 2021; Vento-Tormo et al., 2018). Unexpectedly, both algorithms showed pericytes to be the cell type exerting the highest number of paracrine interactions, most notably with endothelial and glioma cells (Figures 1C, S1B and S1C), despite the low abundance of pericytes (appr. 0.1% of total cells; Figure S1D). Interestingly, whereas glioma cells received signals to a higher extent than sending, pericytes showed a clear dominance in signaling towards other cell types, implying orchestration of distinct niches (Figure 1D). Together, these data demonstrate a vigorous crosstalk between pericytes, glioma cells and the TME, and support the notion of pericytes constituting a signaling hub in glioma.

**Figure 1.**
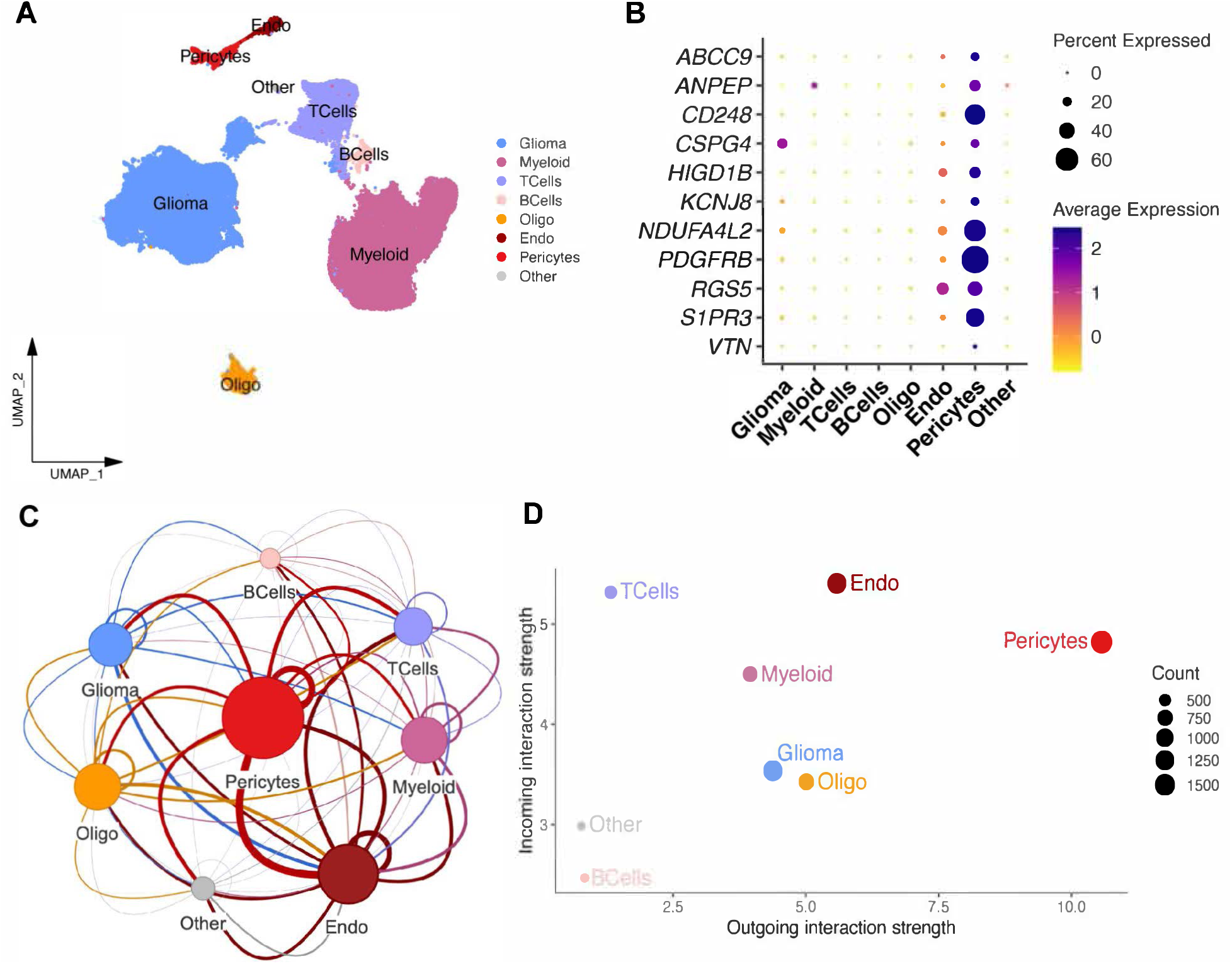
Pericytes form a pivot of intercellular communication in human high-grade glioma. (A) UMAP of human glioma (data from (Abdelfattah et al., 2022), color-coding corresponds to major cell lineages. (B) Average expression of prototypical pericyte marker genes across the cell groups annotated by Abdelfattah et al. (C) A network of human glioma cell clusters, showing the overall cellular communication (autocrine and paracrine), as assessed by CellChat. Node sizes represent the weighted (by the interaction score) total number of interactions, while edges show the weighted number of interactions. (D) Scatterplot showing the total outgoing or incoming communication probability (incoming or outgoing interaction strength) associated with each cell population in the Abdelfattah dataset. Dot sizes represent the number of inferred connections (both outgoing and incoming).

### Pericyte-deprivation leads to an accelerated tumor progression and premature death in two glioma mouse models

To functionally assess the role of pericytes in glioma development and progression, we stereotactically injected *GBM* cells into the subventricular zone (SVZ) of wildtype mice or of mice carrying a homozygous *Pdgfb* allele that is missing the PDGF-B heparan sulfate proteoglycan binding motif sequence (*Pdgfb^ret/ret^* mice) (Lindblom et al., 2003). In these mice, PDGF receptor-β (PDGFRB)-expressing pericytes, which under normal circumstances are recruited to nascent blood vessels through PDGF-BB secretion by endothelial tip cells, are not properly investing the vessel wall due to the increased diffusability of the PDGF-BB ligand in *Pdgfb^ret/ret^* mice (Figure 2A). Hence, a high degree of pericyte loss in the vasculature (70-80%) has been validated in several reports (Abramsson et al., 2003; Andaloussi Mae et al., 2020; Armulik et al., 2010; Lindblom et al., 2003; Villasenor et al., 2017). Intriguingly, following stereotactic engraftment of *p53^-/-^*, *H-Ras* over-expressing neurospheres (Hertwig et al., 2012) into the SVZ, *Pdgfb^ret/ret^* mice exhibited a significantly shorter survival time compared to *Pdgfb^ret/+^*and wildtype animals (Figure 2B). Upon autopsy, the brains of pericyte-poor mice presented with overtly hemorrhagic lesions containing necrotic areas (Figure 2C). In accordance with the literature, we observed an estimated 70-80% reduction of pericyte coverage of tumor vessels upon implantation of syngeneic GL261 cells in *Pdgfb^ret/ret^* mice, compared to control mice (Figures 2D and F). We validated our findings by utilizing RCAS virus-mediated induction of oncogenic *PDGFB* expression and knock-down of *Tp53* in the SVZ of mice expressing the tv⁃a receptor under the *Nestin* promoter (Ntv-a), resulting in tumors that resemble human oligodendroglioma and display a high-grade histology (Ozawa et al., 2014; Squatrito et al., 2010). Accordingly, we crossed *Pdgfb^ret/ret^* mice with Ntv-a mice and induced tumor formation through intracranial injection of virus-producing DF-1 cells. Confirming our previous results, we observed both a dramatically reduced pericyte vessel coverage, and a significantly shorter latency until the *Pdgfb^ret/ret^* mice succumbed to the progressing brain tumors (Figures 2E and 2F). To better understand the biological processes leading to an aggravated tumor progression upon pericyte depletion, we characterized the tumor development more profoundly. To measure hypoxia in the glioma tissue of our mouse model, we quantified HIF2α expression based on glioma section immunostaining, as well as performed a flow cytometry-based analysis of pimonidazole (PIM) adducts in PDGFB-induced brain tumors from cohorts of *Pdgfb^ret/ret^* and *Pdgfb^ret/+^* mice. Regardless of methodology, we did not detect a significant difference in hypoxia in gliomas of pericyte-deprived mice (Figures S2A and S2B). In contrast, a proliferation analysis, based on the expression of KI67, demonstrated a greater accumulation of KI67^+^ cells in PDGFB-induced tumors from *Pdgfb^ret/ret^ mice* compared to *Pdgfb^ret/+^* littermates (Figures 2G and 2H). To learn more about the tumor growth dynamics, we next transplanted luciferase-expressing *p53^-/-^*, *HRAS* over-expressing glioma cells intracranially into *Pdgfb^ret/ret^* and *Pdgfb^ret/+^* animals and monitored the subsequent tumor growth by measuring the total luciferase bioluminescence. In a direct comparison, we found *Pdgfb^ret/ret^* mice to exhibit a higher luciferase activity than the control animals at all time points (Figure 2I). To better estimate differences in the growth kinetics of *Pdgfb^ret/ret^* and *Pdgfb^ret/+^* mice, we fit the data to a non-linear model and observed a higher glioma growth rate in *Pdgfb^ret/ret^* mice over time (Figure 2J), indicating a growth advantage over *Pdgfb^ret/+^* gliomas, in keeping with their higher proliferative rate and faster induction of end-stage disease.

**Figure 2.**
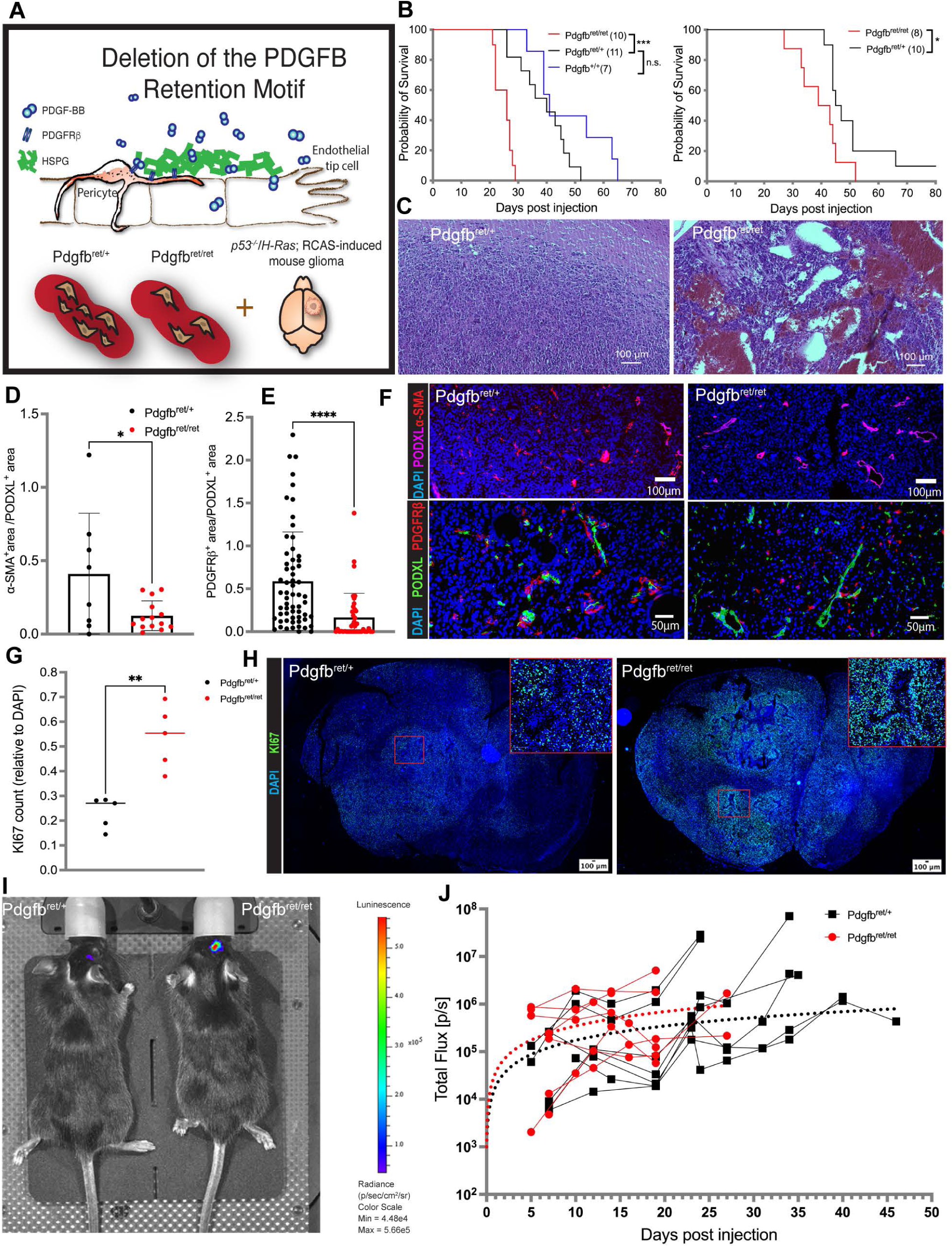
Alterations in glioma development upon pericyte reduction occur in different mouse models. (A) Schematic of tumor cell engraftment or virus mediated glioma generation in *Pdgfb^ret/ret^* mice. (B) Kaplan-Meier curves, showing symptom-free survival of *Pdgfb^ret/ret^*, *Pdgfb^ret/+^*and *Pdgfb^+/+^* mice transplanted with *p53^-/-^*, *H-Ras* over-expressing tumor cells (left panel) and RCAS-PDGFB/shp53 virus producing DF1 cell (right panel). (C) Representative H&E analysis of *p53^-/-^*/*H-Ras* tumor cell induced gliomas in *Pdgfb^ret/ret^* and *Pdgfb^ret/+^* mice. (D-F) Pericyte vessel coverage in GL261 tumor cell (D, upper panels in F) and PDGFB-induced (E, lower panels in F) gliomas in *Pdgfb^ret/ret^* and *Pdgfb^ret/+^* mice, presented as α-SMA^+^/PODXL^+^ and PDGFRβ^+^/PODXL^+^ ratios, derived from the area quantification analysis of immunostained glioma sections (F). (G-H) Proliferation analysis of *Pdgfb^ret/ret^* and *Pdgfb^ret/+^* PDGFB-induced gliomas, presented as KI67^+^ cells relative to DAPI (G), quantified from immunostainings (H). Boxes denote the enlarged regions. (I-J) IVIS imaging of tumor bearing mice from day 5 to 47 after transplantation of luciferase-expressing, *p53^-/-^*/*H-Ras* tumor cells. (I) Representative IVIS scan of a *Pdgfb^ret/ret^* and a *Pdgfb^ret/+^* mouse, 15 days post transplantation. (J) Total photon flux, measured continuously from *Pdgfb^ret/ret^* and *Pdgfb^ret/+^* mice, was fitted to a non-linear model (dotted curves), representing an estimate of the tumor growth kinetics. * p<0.05, ** p<0.01, *** p<0.001, **** p<0.0001, ns not significant

In conclusion, our experimental data reveal that loss of pericytes sustained the engraftment and progression of highly proliferative brain tumors.

### Human GBM heterogeneity is conserved in a *Pdgfb*-driven, *p53^-/-^*murine high-grade glioma model

Based on our observations that pericytes are strong paracrine signal senders in glioma and instruct both tumor and stromal cells, and that their depletion exacerbates tumor progression, we aimed to better understand how pericytes influence the different glioma compartments and their interactions. To that end, we performed scRNA-seq of dissociated tumors from four *Pdgfb^ret/ret^* and seven *Pdgfb^ret/+^* mice, carrying PDGFB-induced glioma tissues. After quality control, we retained a total of 16,086 and 42,838 cells, respectively. We integrated data from both *Pdgfb^ret/ret^*and *Pdgfb^ret/+^* mice together to remove batch effects and to uncover shared cell types between conditions. Unsupervised, graph-based clustering recognized 21 unique clusters (Figure 3A). To annotate clusters, we performed differential gene expression analyses (one cluster *vs* all others) to identify markers that are differentially expressed between clusters but conserved between conditions. We compared significantly over-expressed genes of each cluster to literature-based cell markers (Chanoch-Myers et al., 2022; Couturier et al., 2020; Darmanis et al., 2017; Eberhart and Bar, 2020; He et al., 2016; Mizrak et al., 2019; Neftel et al., 2019; Ochocka et al., 2021), and analyzed the association of GO terms (Metascape) with the top 50 differentially expressed genes (DEG) of each cluster (Zhou et al., 2019) (Figure 3B, Table S1). Tumor cells were identified based on the expression of an RCAS virus-unique gene sequence (Figure 3C). Our analyses revealed four main groups, namely immune clusters annotated as microglia, macrophages, and NK/T/NKT cells (I); a tumor cell-enriched group representing cells expressing marker genes for the astrocytic, oligodendrocytic or neuronal lineages, and tumor cells of hypoxic and mesenchymal character (II); dividing cells (mostly malignant) (III), and vascular cells (IV). Notably, we found only minor fractions of the stromal clusters and a cluster of predominantly physiologic oligodendrocytes to express the RCAS-unique gene sequence. We interpreted these RCAS-positive cells to represent either stromal or differentiated neural cells that were infected by virus particles due to an active *Nestin* promoter, or tumor cells with a high differentiation potential that could take on a non-tumoral phenotype.

**Figure 3.**
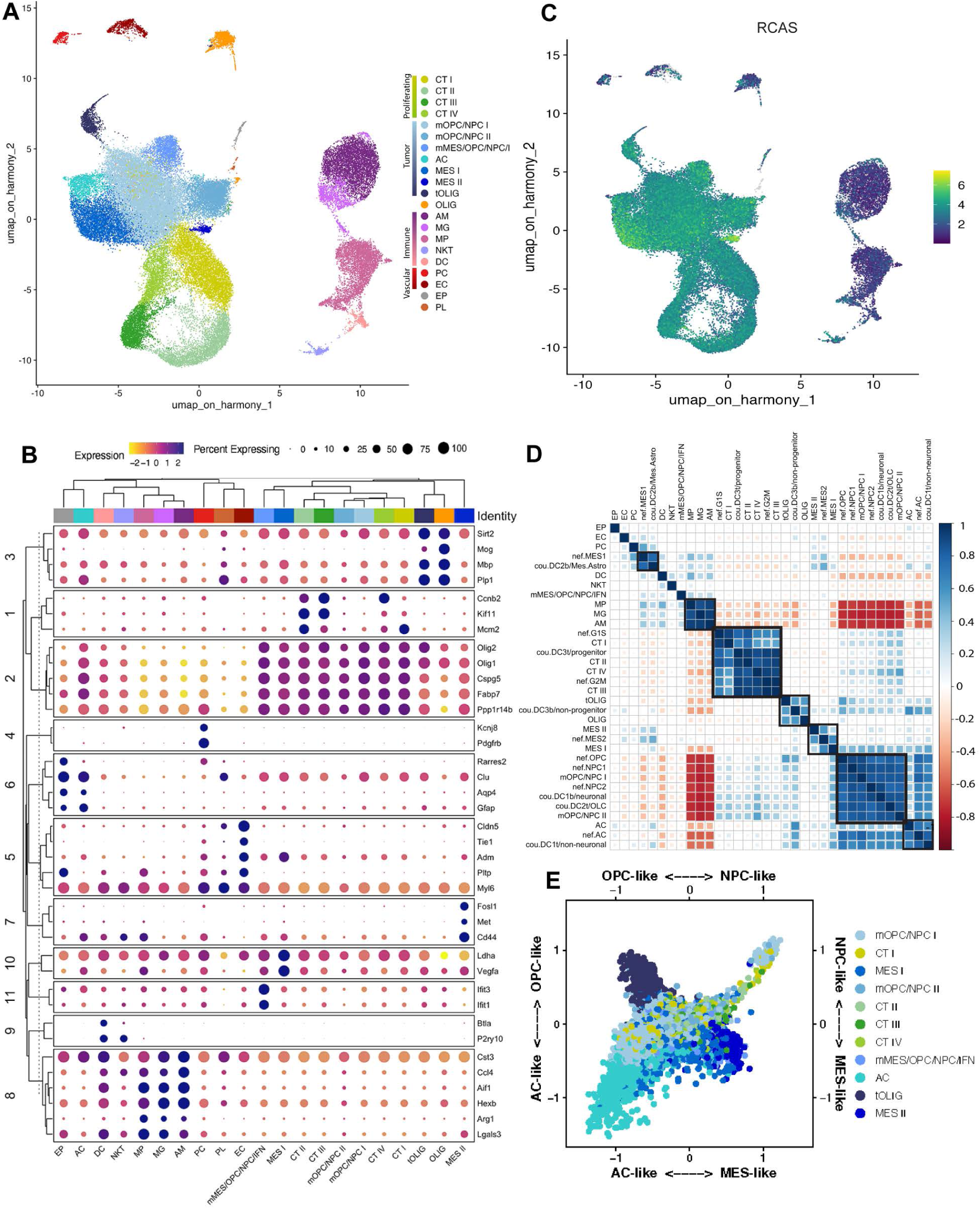
Molecular characteristics of glioma tumor and stromal cells derived from a pericyte-deprived mouse model. (A) UMAP plot of 16,086 *Pdgfb^ret/ret^* and 42,838 *Pdgfb^ret^*^/+^ cells that were isolated from a total of 4 and 7 tumors respectively, and analyzed with scRNA-seq. The UMAP clusters are colored based on cell identify as cycling tumor cells I-IV (CT I-IV), mixed OPC-like/NPC-like tumor cells I-II (mOPC/NPC I-II), mixed mesenchymal/OPC-like/NPC-like/IFN tumor cells (mMES/OPC/NPC/IFN), astrocytes/AC-like tumor cells (AC), mesenchymal tumor cells I-II (MES I-II), tumor oligodendrocytes (tOLIG), oligodendrocytes (OLIG), activated microglia (AM), microglia (MG), macrophages (MP), NK/T/NKT cells (NKT), dendritic cells (DC), perivascular cells/pericytes (PC), endothelial cells (EC), ependymal cells (EP) and platelets (PL). (B) Clustered dot plot showing the scaled average expression of selected top differentially expressed genes across all clusters. (C) Feature plot showing the log-normalized expression of the RCAS-unique vector sequence across all clusters. (D) Clustered correlation matrix showing pairwise Pearson’s correlation coefficients of the mean Z-score estimated for different cell identity signatures from this study, Neftel et al. (nef) and Couturier et al. (cou). Black boxes highlight selected clustered signatures. (E) Butterfly plot of molecular subtype signature scores defined by Neftel et al. The quadrants correspond to the four subtypes: mesenchymal-like (MES-like), neural-progenitor-like (NPC-like), astrocyte-like (AC-like) and oligodendrocyte-progenitor-like (OPC-like). The position of each cell reflects its relative signature score across both axes. Colors represent different clusters as in panel A.

The fact that cluster mOPC/NPC I contributed 25% of all analyzed cells, and the observation of a large proportion of dividing OPC-like cells (Figure S3A) among the analyzed samples identified our murine tumor system as predominantly representing the proneural molecular subtype, which is in line with previous findings; the Ntv-a;RCAS-*PDGFB* murine glioma model has been described to exhibit features of human oligodendrogliomas and proneural GBM subtypes respectively, with *Olig2*-expressing glioma cells constituting the bulk of the tumor (Couturier et al., 2020; Ozawa et al., 2014).

Neftel *et al*. have extended the concept of intra-tumoral heterogeneity in glioma by describing the co-existence of four different states of human GBM cells that recapitulate neural-progenitor (NPC-like) or oligodendrocyte-progenitor (OPC-like) cell states, resemble astrocytes (AC-like), or represent a complex, mesenchymal-like (MES-like) phenotype (Neftel et al., 2019); findings that were corroborated by Couturier *et al*. (Couturier et al., 2020). To test whether our mouse model reflects these cell states that are commonly observed in human patients, we used the gene signatures provided by Neftel et al. and Couturier et al. (Table S2). In short, for each phenotypic gene signature, we calculated the mean z-score for the individual cells in our dataset. We did the same for our own cluster signatures. Then, we computed a pairwise correlation matrix between all signature scores and applied hierarchical clustering to group signatures, based on how similarly they were expressed across our dataset (Figure 3D). The clustering of the tumor and dividing cell signatures from Neftel *et al*. with our tumor clusters’ signatures proved the intra-tumoral heterogeneity and proliferative capacity of the tumor samples (Figures 3D, 3E and S3A). Notably, the MES2 signature grouped with our clusters MES I and II, and the AC-like and “non-neural” signatures from Neftel *et al*. and Couturier *et al*. associated with our cluster AC (Figures 3D and S3A). Except for a tendency towards a more pronounced NPC-like tumor compartment (Figure S3B), we did not observe any major deviation regarding the Neftel cell states in *Pdgfb^ret/ret^* as compared to *Pdgfb^ret/+^* tumors (Figure S3C), reflecting the resilience of the dominant OPC/NPC state in our mouse model, and indicating that the phenotypical aberrance that we observed in glioma development upon pericyte removal was due to more subtle alterations in glioma biology.

### Pericyte deprivation remodels the glioma perivascular niche and impacts on the immune cell landscape

Based on our findings that pericytes are extensively communicating with both tumor-and non-tumor cells, we anticipated an altered glioma microenvironment to contribute to the premature death of pericyte-deprived mice, and therefore focused our attention on the tumor stroma, commencing with the PVN. Pericytes/perivascular cells and endothelial cells could be identified in our dataset based on the expression of marker genes such as *Abcc9*, *Kcnj8*, *Rgs5*, and *Cldn5*, *Tie1*, and *Pdgfb*, respectively (Figure 3B). Indeed, in keeping with human tumors, when we analyzed cell-cell communication networks we found pericytes to exert the most paracrine interactions in both *Pdgfb^ret/ret^*and *Pdgfb^ret/+^* tumors (Figures S4A and S4B).

We next wanted to understand if structural changes occurred upon the deprivation of pericytes. To further validate the *Pdgfb^ret/ret^*mouse model, we quantified the number of cells comprising the PC and EC clusters, and found, as anticipated, the amount of pericytes to be reduced by 51% in *Pdgfb^ret/ret^* mice (Table S3). In sharp contrast, the EC population was increased by 40% (Table S3), in line with previous findings from investigations on brain angiogenesis (Hellstrom et al., 2001). Moreover, measurements based on the labeling of the endothelium with antibodies directed against Podocalyxin provided evidence of structural differences between the *Pdgfb^ret/ret^* and *Pdgfb^ret/+^* glioma vasculature, as we found vessel area, vessel length and number of junctions to be significantly higher in pericyte-poor tumor tissue (Figures 4A and 4B). Finally, we performed a functional analysis on the intactness of the blood-brain barrier by perfusion with labeled dextrans (70kDa) that indicated vascular fluids to leak into the perivascular space (Figure 4C), revealing a significant loss of vascular integrity upon pericyte reduction. The findings of a corrupted endothelium due to pericyte deprivation raised the question whether a structurally altered PVN would have consequences for glioma progression. Since the concept of tumor cell migration along endothelial routes in a vessel co-opting manner is well established (Gupta et al., 2024; Holash et al., 1999; Seano and Jain, 2020), we next aimed at quantifying this process in *Pdgfb^ret/ret^*and *Pdgfb^ret/+^* gliomas. Based on the observation that the vast majority of glioma cells in our mouse model is of OPC/NPC character, expressing *Olig2*, and that these cells tend to invade the brain by single-cell vessel-cooption (Griveau et al., 2018), we performed an immunostaining analysis and quantified the number of OLIG2^+^ cells related to endothelial cells at the invasive rim of mouse tumor-brain tissue sections. We observed two different invasion modes of OLIG2^+^ tumor cells, namely the diffuse spread of individual cells into the surrounding physiological brain parenchyma, and vessel co-option of single cells. Interestingly, we did not detect differences regarding the total number of invading OLIG2^+^ tumor cells in the invasive zone but found a significantly higher proportion of vessel co-opting tumor cells in *Pdgfb^ret/ret^* animals (Figures 4D, 4E, S4C and S4D), indicative of a more permissive PVN in the absence of pericytes.

**Figure 4.**
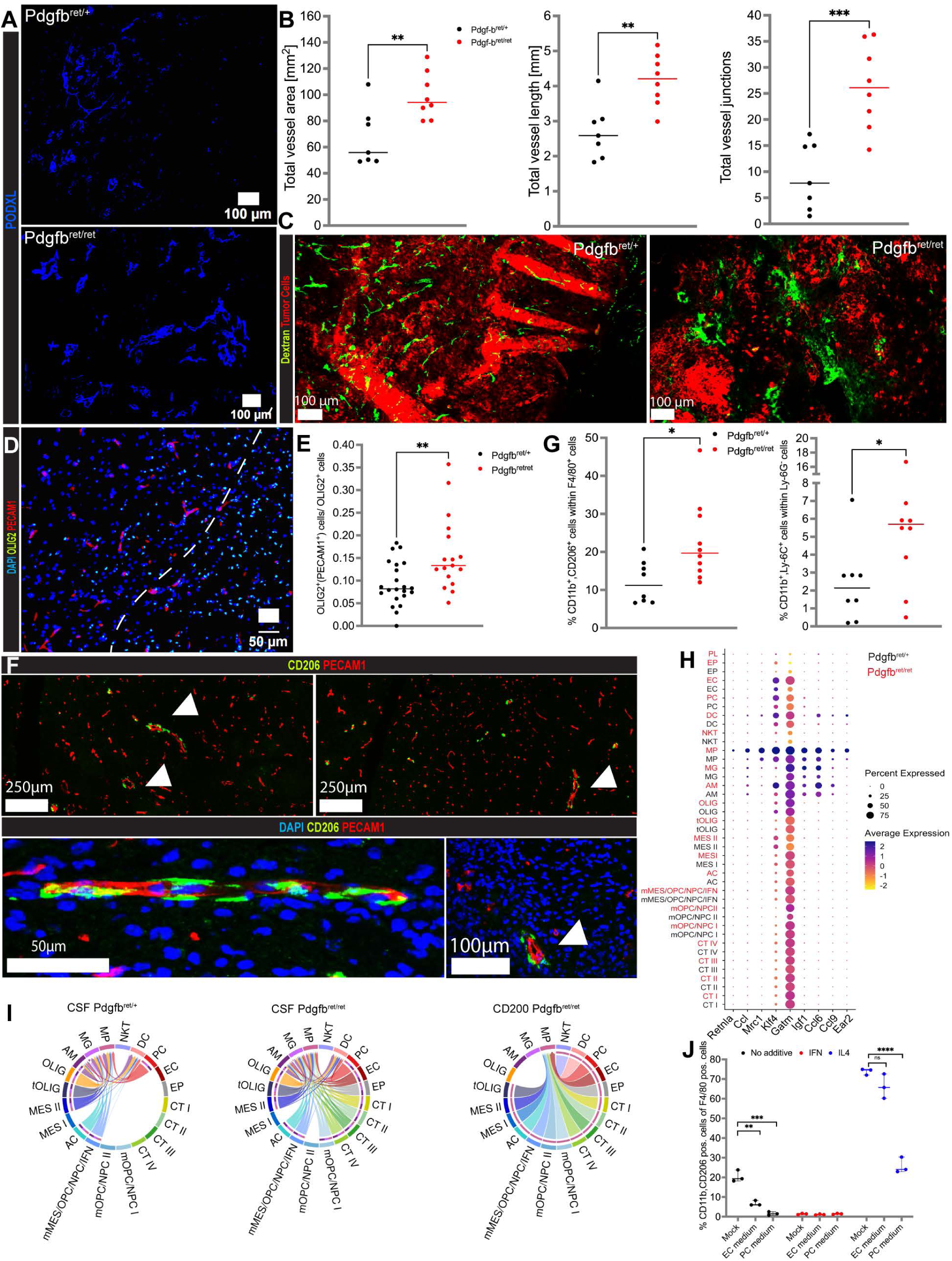
Pericyte reduction causes remodeling of the perivascular and immune glioma microenvironment. (A) Representative immunostainings of PODXL in *Pdgfb^ret/ret^* and *Pdgfb^ret/+^ p53^-/-^*/*H-Ras* tumor cell induced glioma sections. (B) Assessment of glioma tissue parameters total vessel area, length and junctions, comparing *Pdgfb^ret/ret^* and *Pdgfb^ret/+^* tissue sections of *p53^-/-^*/*H-Ras* tumor cell induced gliomas. The measurements were done based on immunostaining of CD31. (C) Representative fluorescence images of a blood vessel functional analysis comparing *p53^-/-^*/*H-Ras* tumor cell induced glioma tissue of *Pdgfb^ret/ret^* and *Pdgfb^ret/+^* mice. Terminally sick animals were perfused with labeled dextran prior to euthanizing. (D-E) Quantification of tumor cell infiltration at the invasive rim (IR) in PDGFB-induced gliomas derived from *Pdgfb^ret/ret^* and *Pdgfb^ret/+^*mice. Measurements were performed based on immunostaining of PECAM1 and OLIG2. In an approximation, the IR was defined as the field of view (FOV) portion adjacent to the cell dense tumor core (white dotted line) (D). Vessel co-opting tumor cells were defined as OLIG2^+^ cells in contact with PECAM^+^ cells and normalized to total OLIG2^+^ cells (E). (F) Representative images of vessel-associated macrophages showing immunostainings of CD206 and PECAM1. Arrow heads indicate CD206^+^ cells adjacent to PECAM1^+^ cells. (G) FACS-based quantification of CD206^+^/CD11B^+^/F4/80^+^ macrophages (left panel) and CD11B^+^/LY6C^+^/LY6G^-^ myeloid-derived suppressor cells (MDSC, right panel), comparing PDGFB-induced *Pdgfb^ret/ret^*and *Pdgfb^ret/+^* glioma tissue. (H) Dot plot showing the scaled average expression of selected macrophage polarization markers in the scRNA-seq data across all clusters, comparing *Pdgfb^ret/ret^* and *Pdgfb^ret/+^* cells. (I) Chord diagrams of selected identified signaling pathways that exhibit different signaling patterns between *Pdgfb^ret/ret^* and *Pdgfb^ret/+^* tumors, performed with CellChat. In the chord plots, the edge weights are proportional to the estimated interaction strength. Left and middle panel: CSF pathway (active in both *Pdgfb^ret/ret^* and *Pdgfb^ret/+^*tumors); right panel: CD200 pathway (active exclusively in *Pdgfb^ret/ret^*tumors). (J) FACS-based quantification of CD206^+^ murine bone marrow derived, proliferating macrophages, cultured with pericyte-or endothelial cell-primed or mock medium. Either IL4, IFNγ or no cytokine was added to the cultures. Analysis was performed 48h post addition of primed media. * p<0.05, ** p<0.01, *** p<0.001, **** p<0.0001, ns not significant

Next, we set out to investigate putative alterations of the immune cell landscape in pericyte-poor tumors. Since an accumulation of myeloid cells with tumor-supportive features has been shown for experimental pericyte-poor tumors (Hong et al., 2015), we focused on the prevalence of immune-suppressive, polarized macrophages that are typically characterized by the expression -among others-of *Cd206*, *Cd204*, *Arg1*, and *Lgals3* (Pombo Antunes et al., 2021). In tissue sections of tumors from both *Pdgfb^ret/ret^* and *Pdgfb^ret/+^* mice, we found CD206^+^ macrophages to be exclusively associated to blood vessels (Figures 4F), indicating a preference for, and/or an induction of a polarized phenotype by the perivascular space . Using flow cytometry, we quantified two populations of alternatively activated myeloid cells that have been described as immune-suppressive, defined by positivity for CD206, CD11b, and F4/80 (polarized macrophages), as well as the marker expression combination CD11b^+^/LY6C^+^/LY6G^-^(myeloid-derived suppressor cells, MDSC), and observed a significantly higher cell number in *Pdgfb^ret/ret^* tumors for both cell populations (Figure 4G). Intriguingly, we also noted a shift towards a more extreme polarization for macrophages in pericyte-deprived tumors, as these cells exhibited a higher expression of genes associated with alternative activation such as *Retnla*, *Ccl24*, *Mrc1* (CD206), *Arg1*, and *Klf4* (Figure 4H, Table S4). Whereas an increased accumulation of myeloid cells could, at least partially, be explained by a more permeable vasculature, this structural alteration is unlikely to impact on the character of macrophages. Therefore, we scrutinized the signaling profile of the endothelium in our scRNA-seq data with CellChat. Of note, three signaling axes known to induce macrophage polarization -CD200, CSF, and GAS6 (Jeannin et al., 2018; Sousa et al., 2015; Wu et al., 2018) -were exclusively active in *Pdgfb^ret/ret^* endothelial cells that signaled towards macrophages (Figures 4I and S4E), and presumably instructed them toward an immune-suppressive phenotype.

Lastly, we investigated whether pericytes might directly exert a dampening, anti-polarizing effect on myeloid cells. To this end, we applied supernatant collected from mouse brain pericyte and endothelial cell cultures to proliferating macrophages and included IFNγ-“non-polarized” and IL4-“super-polarized” controls. Strikingly, pericyte supernatant, in contrast to endothelial cell supernatant, dampened the IL4-related “super-polarization” effect four-fold, indicating a putative direct paracrine, polarization-suppressing effect of pericytes (Figure 4J).

In summary, our results show that the perivascular loss of mediation due to deprivation of pericytes leads to remodeling of the perivascular niche, characterized by dysregulated vessel growth, reduced patency of the blood-brain barrier, increased tumor cell vessel co-option, altered endothelial cell signaling, and an abundance of polarized immune-suppressive macrophages.

### A hypoxia-associated subset of GBM cells harbors a distinct mesenchymal state

Since previous work has shown that the MES-like GBM cell state is not only correlated to the abundance of macrophages, but directly driven by alternatively activated myeloid cells (Chanoch-Myers et al., 2022; Hara et al., 2021; Sorensen et al., 2018; Wang et al., 2018), glioma features that we too had observed in our study, we scrutinized the MES I and II cells of our scRNA-seq dataset more thoroughly. Among the 50 most highly expressed genes in MES I, we found many to be related to hypoxia induction and regulation, such as *Vegfa*, *Ldha*, and *Pgk1* (Figure 5A). Conversely, MES II featured a completely different set of top DEGs. Whereas we again identified hypoxia and stress-related genes like *Hmox*, *Tnc*, *Gdf15*, and *Hspa9*, MES II marker genes also included *Fosl1*, *Met*, *Ccn4*, *Grb10,* and *Sema3c*, genes that have previously been associated with stemness, tumor cell plasticity and invasion, the regulation of EMT-like processes, and generally an increased tumor aggressiveness (Chen et al., 2022; Hao et al., 2023; Marques et al., 2021; Qin et al., 2020; Tao et al., 2020). Moreover, a transcription factor analysis with ProgenY (Schubert et al., 2018) showed a similar activity pattern between MES I and MES II cells but indicated a much higher transcription factor activity for MES II cells (Figure S5A). Interestingly, several MES II marker genes such as *Ccn4*, *Tnc*, and *Grb10* were co-expressed by pericytes, cells of mesenchymal character, sometimes also considered as mesenchymal stem cells (Armulik et al., 2011). Thus, whereas the expression profiles of both MES I and II cells appeared to be related to hypoxia, only the profile of MES II cells represented a panel of distinct mesenchymal features. Notably, the overall percentage of tumor cells decreased, whereas we observed the opposite trend for MES II cells when comparing *Pdgfb^ret/+^* versus *Pdgfb^ret/ret^* gliomas (34% increase in *Pdgfb^ret/ret^* mice; Table S3 and Figure S3C), indicating that the loss of pericytes favors the expansion of this extreme mesenchymal glioma cell state. This notion was further supported by cell-cell communication analysis that demonstrated an increased signaling activity of pathways related to mesenchymal transformation, such as Wnt, Notch, and Gas6/Axl (Bazzoni and Bentivegna, 2019; Sadahiro et al., 2018; Zhan et al., 2017) in MES II cells of *Pdgfb^ret/ret^* samples (Figures S4E and S5B).

**Figure 5.**
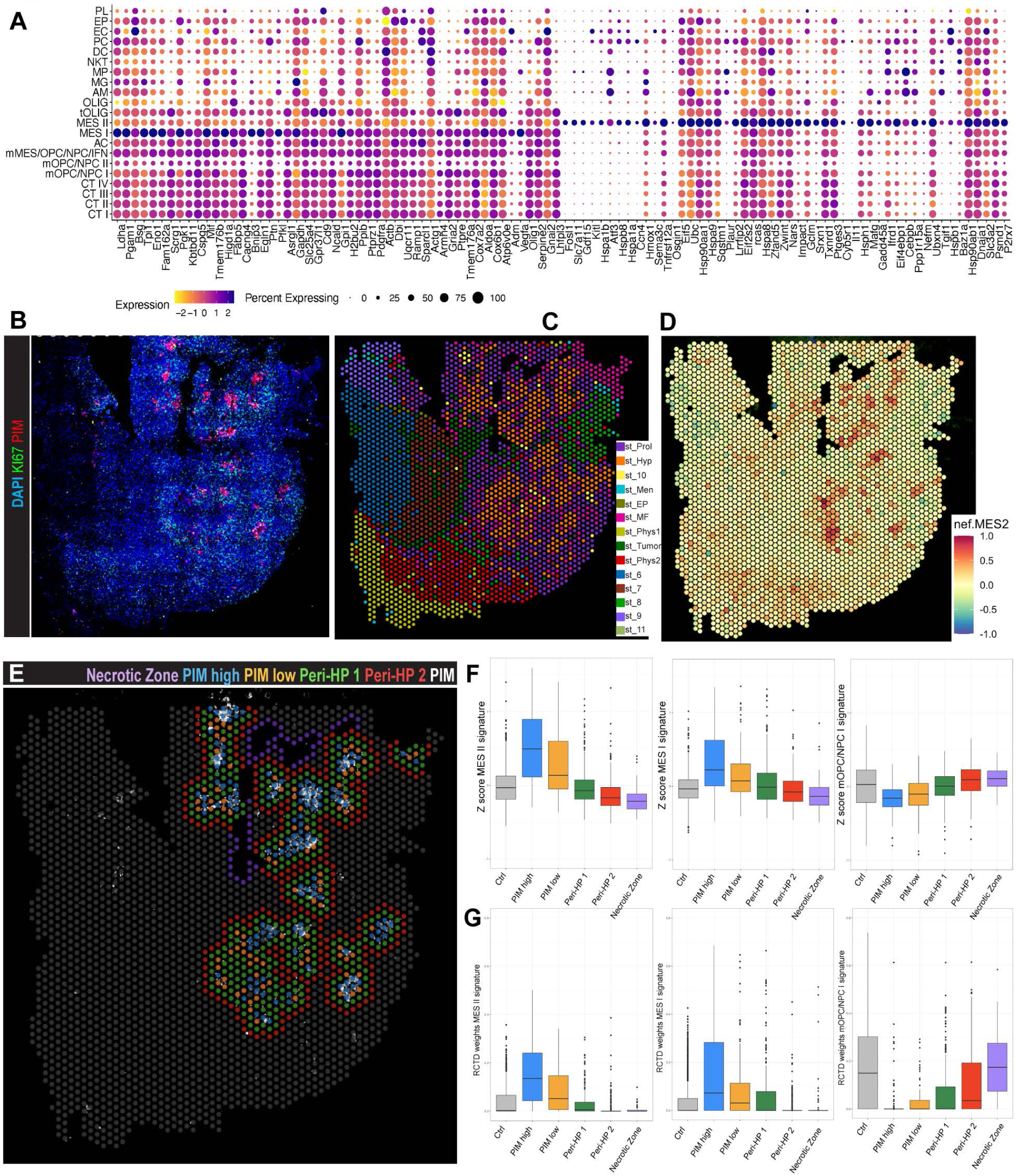
Localization and functional characterization of mesenchymal cell subgroups with spatial transcriptomics analysis. (A) Dot plot showing the scaled expression of the MES I and MES II top 50 differentially expressed genes (combined *Pdgfb^ret/ret^* and *Pdgfb^ret^*^/+^) across all scRNA-seq clusters. (B) Representative image of a *Pdgfb^ret/ret^* glioma section (stRNA-seq sample #1159) mounted onto the capture area of a Visium Gene Expression Slide (VGES), and immunostained against PIM and KI67. (C) Image of a *Pdgfb^ret/ret^* glioma section (#1159), mounted on a VGES capture area. 14 spatial clusters were annotated based on the integration of all 8 samples (four *Pdgfb^ret/ret^*and four *Pdgfb^ret^*^/+^ glioma sections) using Harmony. (D) Image of a *Pdgfb^ret/ret^*glioma section (#1159) colored by the mean Z-score of the MES2 signature from Neftel et al. (E) VGES capture area of stRNA-seq sample #1159, showing a manual annotation of a hypoxic gradient. Hypoxic zones are highlighted by PIM positivity ("PIM high" and "PIM low"), while peripheral zones are PIM negative ("Peri-HP1" and "Peri-HP2"). (F-G) Boxplots showing the prevalence of MES II and mOPC/NPC I cells by the average Z-score (F) and RCTD proportions (G) per spot in manually annotated hypoxic and peri-hypoxic zones defined in panel (E). All remaining capture spots served as control zone.

To interrogate if MES II cells were more prevalent in certain tumor biomes (Barthel et al., 2022), we explored our glioma samples in a spatial context using array-based, spatial transcriptomics RNA-sequencing (stRNA-seq) technology (10x Genomics Visium), and simultaneously visualized blood vessels, hypoxic regions, and proliferative tissue areas by immunostaining of Podocalyxin (PODXL), PIM adducts, and KI67 (Figures 5B and S5C). To spatially characterize the tumor sections, we integrated the data of four *Pdgfb^ret/ret^* and four *Pdgfb^ret/+^* tumor sections using Harmony (Korsunsky et al., 2019), and overlaid the transcriptomics data with the immunostaining images. The expression profile of each Visium spot is a mixture of the transcriptome of several cells that are likely of different cell types. Therefore, these clusters represent niches or tumor zones, rather than cell types. To annotate these clusters, we considered IF stainings, DEGs, and estimated scRNA-seq cluster z-scores. We annotated 14 spatial clusters of predominantly tumor or physiologic phenotype (Figures 5C and S5D). Whereas we found cluster st_Hyp to overlap with hypoxic, PIM^+^ areas, and to feature a generally hypoxic tumor cell expression signature, these zones were surrounded by KI67^+^ st_Prol spots of a dominant proliferation profile that highly correlated with our scRNA-seq proliferation signatures of clusters CT I-IV (Figures 5C, S5E and S5F). Interestingly, st_Hyp-dominated tissue areas showed the strongest expression of the Neftel *et al*. MES2 and our MES II signatures (Figures 5B, 5D and S5F). Moreover, macrophage polarization marker *Msr1* (CD204) appeared to be mostly expressed in hypoxic zones (Figure S5G). Of note, we observed st_MF capture spots, that had a high expression of genes typical for myofibroblasts (*Col1a1*, *Col1a2*, *Lum*, *Dcn*; Table S5), interspersed among st_Prol and st_Hyp capture spots (Figures 5C and S5D). Furthermore, we found st_Tumor to be spatially separated from st_Hyp, st_Prol and st_MF and to feature a mixed AC, mOPC/NPC I-like expression profile (Figures 5C, S5E and S5F). The remaining spatial clusters were either annotated based on their expression of genes that are functionally related to neuronal processes and considered as glioma cell invasion zones (st_Phys1 and st_Phys2), or their spatial occurrence at meningeal zones and brain cavities lined by epithelial cells (st_Men and st_EP), or could not be specified (st_6-11) (Figures 5C, S5E and S5F).

A hierarchical clustering analysis of the integrated spatial clusters corroborated our annotations, showing that the Visium capture spots are dominated by either a glioma/hypoxia/myofibroblast or a physiologic neural/meningeal expression profile (Figure S5H). Moreover, to provide further support for the spatial cluster annotation, and to characterize our samples on a deeper level, we estimated the cell type composition of spots by applying the Robust Cell Type Decomposition (RCTD) method (Cable et al., 2022). RCTD infers the most likely proportion of different cell types in every Visium spot, by using a scRNA-seq dataset as reference. Here, we used the expression profiles from our 21 scRNA-seq clusters as reference and found, in almost perfect congruence with the mean Z-score estimations, RCTD to show a zonation of the tumor samples into a MES, an OPC/NPC, a proliferative and a physiologic invasive zone, with the MES and the OPC/NPC sector being mutual exclusive (Figure S5E).

Notably, when we quantified the abundance of the MES II signature in a Z-score-and RCTD-based analysis for the st_Hyp clusters of the spatial dataset, both methods indicated an increase in MES II cells in *Pdgfb^ret/ret^* as compared to *Pdgfb^ret/+^* samples (Figure S5I), which is in line with our observations from the scRNA-seq dataset. To determine the prevalence of MES II cells, we manually annotated the core hypoxic, as well as the peri-hypoxic zones of the spatial glioma sections, and visualized the distribution of MES II signature mean z-scores and RCTD proportions across spots (Figures 5E, 5F and 5G). We observed the highest expression of MES II signature z-scores and the highest prevalence of MES II cells in the hypoxic core regions, and a gradual decrease of the expression intensity and prevalence towards the hypoxic periphery, while mOPC/NPC I cells exhibited a reverse distribution pattern.

Whereas this result confirmed hypoxic gene expression as a central element of the MES II cells, we wanted to find out more about the MES II signature genes that were functionally related to other mesenchymal glioma processes. To this end, we first asked whether there are differences in the spatial expression distribution of the scRNA-seq signatures of MES I and MES II in our stRNA-seq data (Figure 6A and S5E). Of note, whereas MES I expression was restricted to hypoxic zones, MES II expression also appeared at the tumor invasive zone of the tissue section where tumor cells infiltrate physiologic areas of the brain parenchyma; a pattern that could be observed in most *Pdgfb^ret/+^* samples, but even more conspicuous in GBM from *Pdgfb^ret/ret^* mice (Figure 6A). Additionally, the overall expression of the MES II signature was more pronounced in the absence of pericytes, in keeping with the higher abundance of these cells observed in the scRNA-seq analysis (Figure 6A). These findings indicated alternative functions of MES II cells, exceeding a mere hypoxia-induced gene expression program. Interestingly, we found the MES II top marker *Fosl1* to be expressed both in hypoxic and tumor peripheral zones. *Fosl1* is an AP-1 transcription factor, and its expression is correlated to tumor progression and unfavorable prognosis in various epithelial tumor types (Elangovan et al., 2018; Li et al., 2019; Liu et al., 2021; Zhang et al., 2021). It has been found previously to be induced by hypoxia, and represents a master regulator of stemness, EMT, and invasion (El Zarif et al., 2024; Marques et al., 2021; Zhang et al., 2023). Based on this observation, we next sought to explore if a subpopulation of MES II cells developed invasive capabilities and migrated away from the st_Hyp/PIM^+^ zones, and toward the tumor periphery. Therefore, we first curated two MES II sub-signatures comprising genes associated to either hypoxia and stemness (HSS; Table S6) or to invasiveness (ISS; Table S6). To see how unique these sub-signatures were, we checked their distribution among the different annotated clusters of our scRNA-seq dataset (Figure 6B). Whereas we found HSS to be clearly up-regulated in MES II cells only, ISS appeared to be moderately up in both MES II and the PC cluster, the latter representing stromal cells with mesenchymal features. When we investigated the spatial context of the sub-signatures, we observed HSS to be mainly expressed around hypoxic zones, whereas ISS expression appeared mainly in the distant, tumor infiltration zone (Figure 6C). We aimed to corroborate these findings by deconvolving the multicellular capture spots of our stRNA-seq dataset with RCDT, and observed MES I and MES II cells to contribute equally to the expression profiles of hypoxic zone capture spots. Conversely, we found a majority of the tumor invasive zone spots to feature the MES II signature only (Figure 6D).

**Figure 6.**
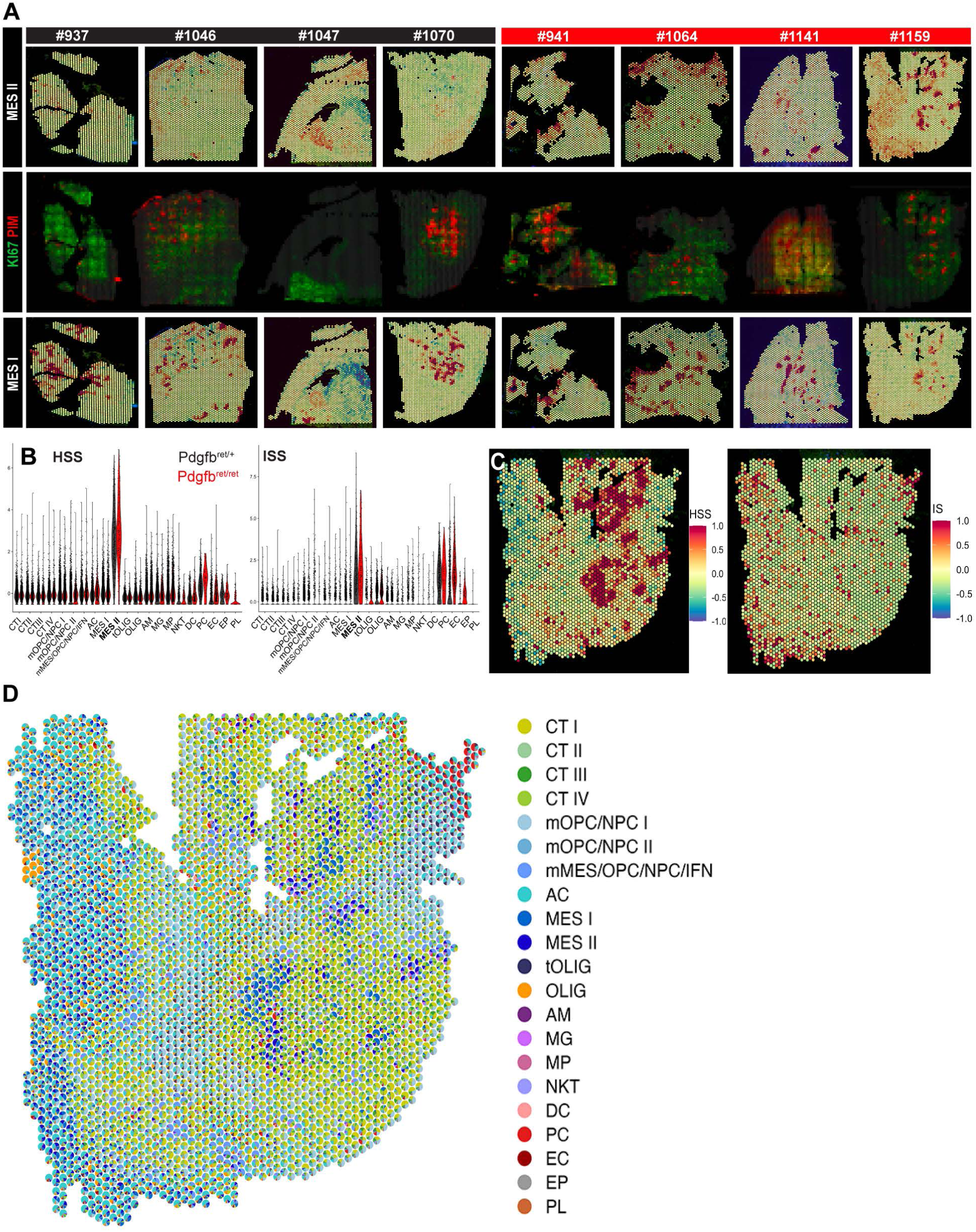
Mesenchymal tumor cells, differing in distinct gene expression programs, prevail in spatially separated glioma niches. (A) VGES capture area images of 4 *Pdgfb^ret/ret^* and 4 *Pdgfb^ret^*^/*+*^ glioma sections (upper and lower panels) colored by the mean z-scores of the scRNA-seq clusters MES I and II. The middle panels show the corresponding immunostainings of PIM and KI67 for each section. (B) Violin plots showing the logNormalized expression of the MES II sub-signatures HSS (hypoxia and stemness) and ISS (invasiveness) across all scRNA-seq clusters for *Pdgfb^ret/ret^* and *Pdgfb^ret/+^*tumor derived cells. (C) Image of a *Pdgfb^ret/ret^*glioma section (#1159) colored by the mean Z-score of HSS (hypoxia and stemness) and ISS. (D) Spatial Scatterpie chart of a *Pdgfb^ret/ret^* glioma section (#1159) showing the RCTD proportions of different celltypes per spot.

In summary, these results underline the unique character of MES II cells, and imply that they are primed in hypoxic regions in contact with polarized macrophages, followed by a development of invasive features and infiltration of the tumor periphery.

### Pericyte depletion reinforces a highly invasive, *Met* expressing subset of mesenchymal GBM cells

Having observed a higher abundance of polarized macrophages in *Pdgfb^ret/ret^*gliomas and their location in mesenchymal tumor areas, we next aimed to investigate the functional dependency between tumor-supportive macrophages and invasive MES II cells in a pericyte-deprived microenvironment. Exploration of our scRNA-seq dataset with CellChat revealed active HGF/MET signaling to occur exclusively between sending, *Hgf-*expressing macrophages and receiving, *Met*-expressing MES II cells in *Pdgfb^ret/ret^*, but not in *Pdgfb^ret/+^* glioma tissue (Figures 7A and 7B). *MET* is frequently over-expressed in human GBM due to amplification, mutation and fusion events (Al-Ghabkari et al., 2024; Cheng and Guo, 2019; Hu et al., 2018; Petterson et al., 2015) and has been demonstrated to promote tumorigenesis through stimulation of glioma cell migration and invasion (Mulcahy et al., 2020). Indeed, when we determined the *Met* expression in our spatial transcriptomics dataset, we found *Met* to be expressed in the same area as the ISS sub-signature (Figure 7C). Notably, we observed a very pronounced invasive pattern with a stRNA-seq sample that consisted of a smaller tumor core and a relatively large physiologic tissue area, including cortical zones: here, *Met* expression was almost completely absent in the *RCAS^+^* tumor core zone, while it could be found in capture spots otherwise dominated by a non-tumorous expression profile (Figures S6A and S6B).

**Figure 7.**
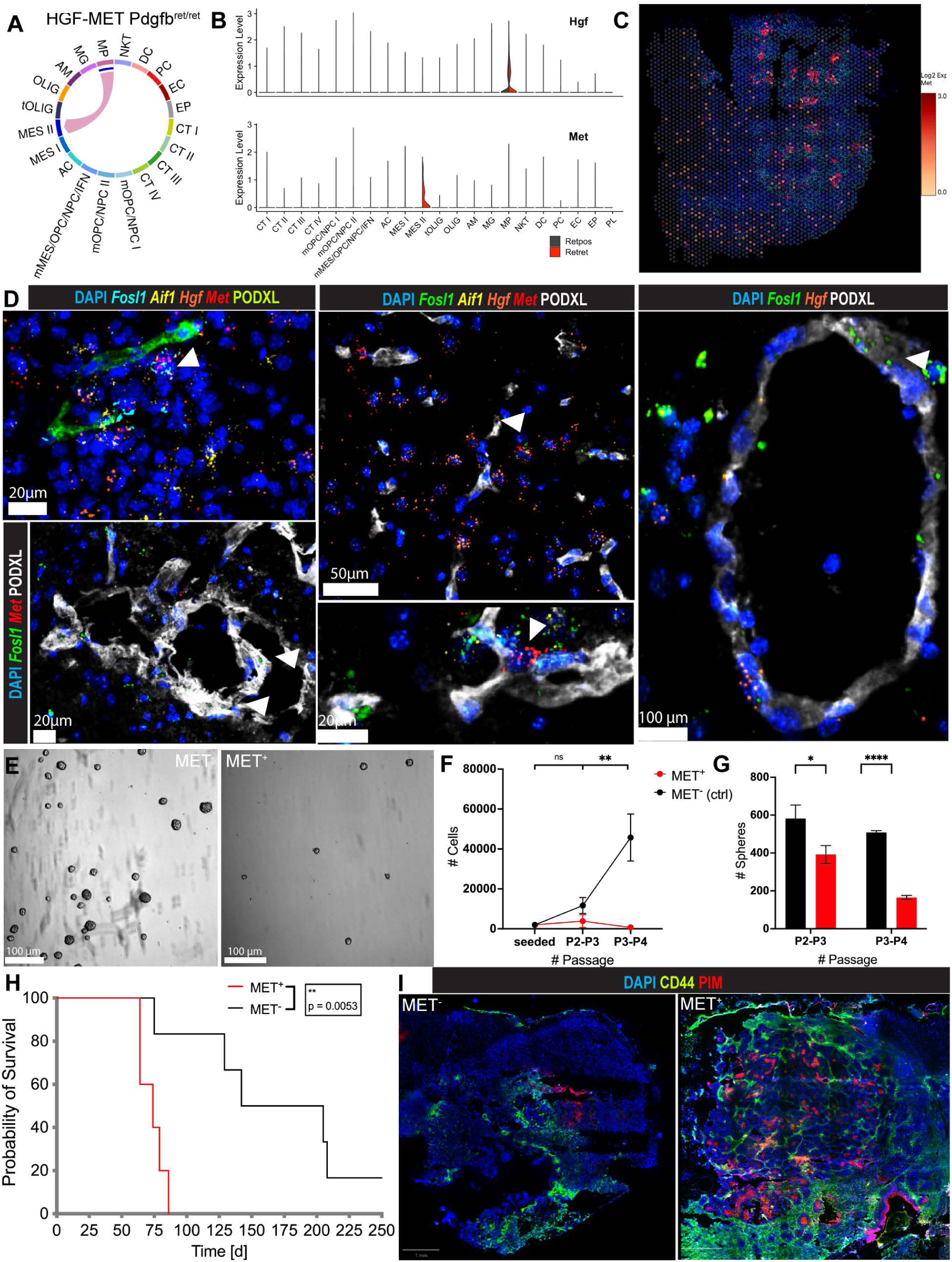
Mesenchymal tumor cells interacting with macrophages drive an invasive glioma phenotype. (A-B) Chord diagram of the HGF-MET signaling pathway, active exclusively in *Pdgfb^ret/ret^*tumors, based on the CellChat analysis. The plot shows the *Pdgfb^ret/ret^*tumor specific interaction of macrophages and MES II cells (A). Violin plot shows the logNormalized expression of *Hgf* and *Met* in *Pdgfb^ret/ret^* and *Pdgfb^ret^*^/*+*^ cells across all scRNA-seq identified clusters (B). (C) Spatial distribution of *Met* expression for a representative *Pdgfb^ret/ret^* stRNA-seq sample (#1159). (D) Combined multiplexed immunostaining-*in situ* hybridization (ISH) analysis for the detection of *Hgf*, *Aif1*, *Fosl1*, *Met* and CD31 in PDGFB-induced gliomas, derived from *Pdgfb^ret/ret^*mice. Arrow heads indicate *Fosl1^+^* and *Fosl1^+^/Met^+^*cells. (E-G) *In vitro* growth parameter analysis of MET^+^, F4/80^-^ and MET^-^, F4/80^-^ cells, isolated from PDGFB-induced glioma tissue. Cells were seeded at passage (P) 2. Phase contrast images were taken at P3 (E) and total cell number (F) as well as total sphere number (G) per culture were determined at passages P3 and P4. (H) Kaplan-Meier curves showing symptom-free survival of mice transplanted with MET^+^ and MET^-^ glioma cells. (I) Representative immunostainings of CD44 and PIM on glioma sections derived from intracranial engraftment of MET^+^ and MET^-^ glioma cells.

To further validate the functional relation between *Met-*expressing cells and macrophages, and to determine their occurrence in distinct glioma niches, we performed a combination of multiplex *in situ* hybridization and immunostaining on *Pdgfb^ret/ret^* glioma sections, using probes against *Hgf*, *Aif1* (IBA-1), *Fosl1*, and *Met* as well as an antibody against CD31. We frequently detected *Hgf^+^/Aif1^+^* cells in the vicinity of *Fosl1* and *Met* expressing cells (Figure 7D). Moreover, we also found cells that co-expressed *Fosl1* and *Met*, indicative of MES II cells (Figures 7D and S6C). Notably, we observed an abundance of *Aif1*, *Met*, and *Fosl1* expressing cells not exclusively in hypoxic regions but also in the perivascular space of *Pdgfb^ret/ret^* glioma sections, indicating a putative opportunistic occupation and exploitation of this glioma niche vacated by pericytes (Figures 7D and S6C).

Due to the reported high tumorigenic and invasive potential of *Met-*expressing cells, and more permissive conditions of the TME upon pericyte depletion, we postulated that MET^+^ cells are among the main drivers of the observed aggravated tumor progression in *Pdgfb^ret/ret^* mice. To investigate this hypothesis, we used flow cytometry to isolate one population of MET^+^, F4/80^-^ and one population of MET^-^, F4/80^-^ cells as a control from glioma tissue and expanded these cells in culture. Interestingly, MET^+^, but not MET^-^ cells proved to be difficult to expand and exhibited both a significantly lower proliferation rate and sphere-induction potential when compared to control cells, implying the dependency of these cells on the glioma microenvironment (Figures 7E, 7F and 7G). In line with this proposition, and in keeping with their mesenchymal phenotype, upon intracranial transplantation, we found MET^+^ cells to generate highly aggressive, 100% penetrant gliomas, that developed significantly faster as compared to transplanted MET^-^ cells (Figure 7H), irrespective of injection into *Pdgfb^ret/+^* or *Pdgfb^ret/ret^*animals (Figure S6D). When we compared tissue sections from MET^+^ and MET^-^ gliomas, we observed a strong expression of the glioma MES-like marker CD44 (Hara et al., 2021), and the abundance of hypoxic areas in MET^+^ tumors, confirming the mesenchymal character of these gliomas (Figure 7I).

Taken together, our studies suggest that pericytes attenuate alternative macrophage activation and interfere with their stimulation of an extreme mesenchymal subpopulation that drives tumorigenesis and glioma cell invasion.

### HSS and ISS expression is associated with poor survival and spatially linked to glioma cell infiltration and the absence of pericytes

In order to find further support in human GBM for a link between the absence of pericytes and a more aggressive GBM phenotype characterized by increased invasion into the surrounding brain parenchyma, we analyzed public patient cohorts with clinical follow-up data and stRNA-seq analysis. First, we assessed the correlation between the MES II signatures derived from our scRNA-seq data with outcome in the TCGA High Grade Glioma (HGG) cohort. Indeed, patients with above-median expression of either MES II HSS, ISS, or the complete MES II signature exhibited a significantly shorter progression-free interval, as well as overall survival (Figures 8A and S7A). Intriguingly, a core pericyte gene expression signature (van Splunder et al., 2024) was expressed at a lower level in recurrent samples from the cohort, compared to the matched primary tumors, in keeping with previous findings that recurrent GBMs exhibit a more aggressive growth pattern with a more pronounced MES signature (Hoogstrate et al., 2023; Wang et al., 2022) (Figure 8B). Next, we aimed to determine the spatial localization of the MES II signature in human GBM samples. To that end, we explored an atlas of spatially resolved transcriptomics, consisting of specimens derived from 20 GBM patients, that were histologically classified applying the Ivy-gap database classification system (Ravi et al., 2022) (Figure S7B). In accordance with observations from our HGG mouse model, a module score analysis revealed a higher expression of the MES II/ HSS genes in mesenchymal tumor zones, characterized by high MES 2 signature expression ("vascular") (Figures 8C, S7B, S7C and S7D). Conversely, the MES II/ ISS genes appeared to be higher expressed in zones that were histopathologically characterized as "infiltrative" and "white matter" (Figures 8C, S7A, and S7E). Finally, we used the core pericyte marker gene signature as a proxy for pericyte abundance, and found it to be upregulated in the "cellular" tumor zone (Figure 8D). In contrast, lower pericyte marker gene expression was observed in the "vascular"/MES 2 high tumor zone, and was virtually absent in the infiltrative tumor areas, indicating that the abundance of pericytes and HSS/ISS-expressing mesenchymal glioma cells are anti-correlated (Figure 8D).

**Figure 8.**
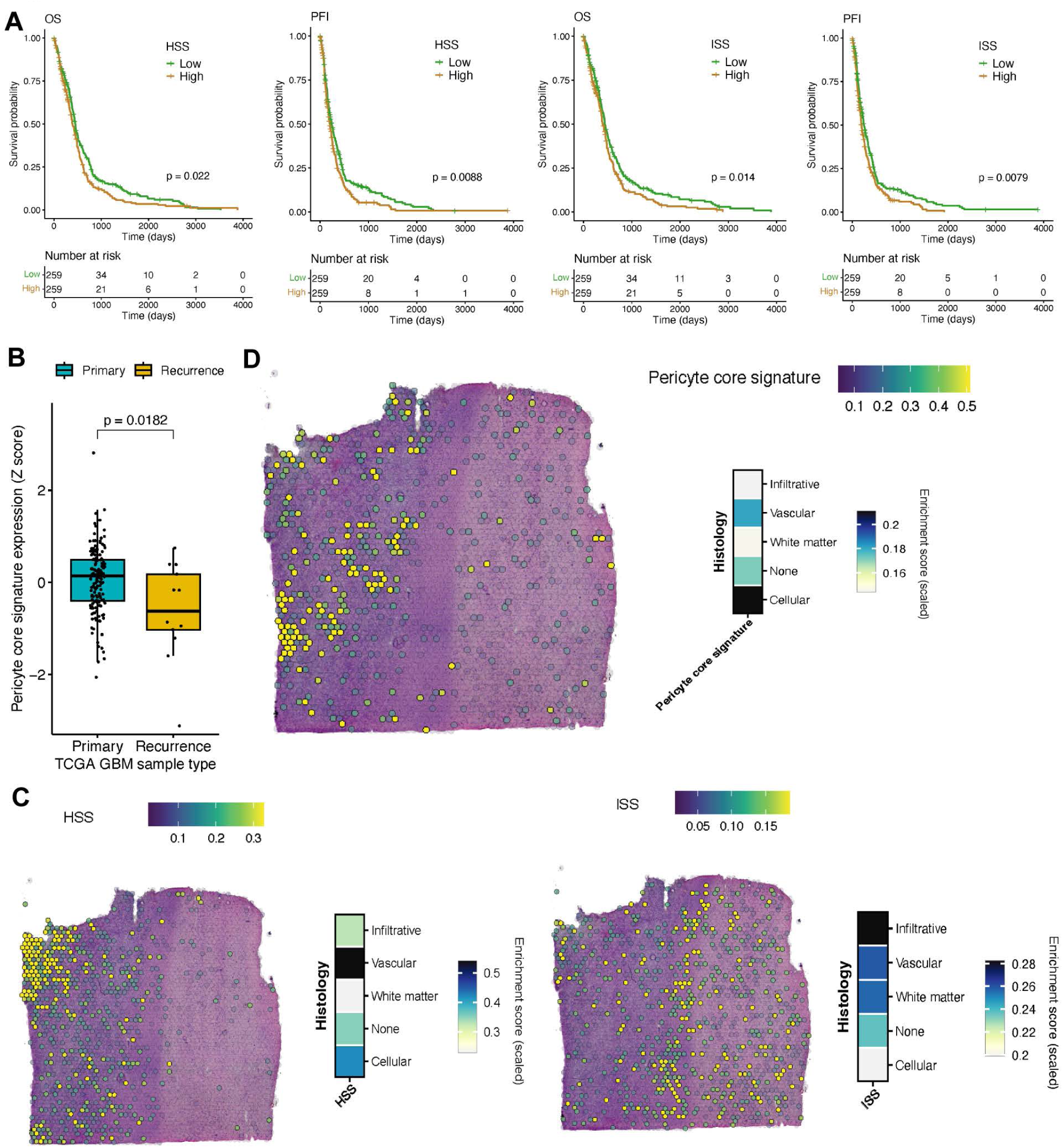
Survival and spatial localization analysis of GBM patient datasets. (A) The Kaplan-Meier curves show the overall survival (OS) and progression-free interval (PFI) probabilities of the high (brown) and low (green) HSS and ISS expression groups in the TCGA GBM cohort (n = 518). p-value: log-rank test. (B) Box plot showing the expression of the pericyte core signature in primary (n = 153) and recurrent (n = 13) TCGA GBM samples. p-value: Wilcoxon test. (C-D) Spatial plots and heatmaps showing HSS (C, left) and ISS (C, right) and Pericyte core gene (D) module scoring in a spatially resolved transcriptomics Human GBM specimen.

Collectively, we propose a model in which the relative absence of pericytes in recurrent human GBM instigates immune-suppressive macrophages to support a pattern of perivascular invasion of glioma cells with extreme mesenchymal features, leading to reduced progression-free and overall survival.

## Discussion

The perivascular space has been elaborately characterized as a microenvironment that drives progression and invasion of human gliomas (Charles and Holland, 2010; Hambardzumyan and Bergers, 2015; Jung et al., 2021). Here, we provide evidence that cells embedded within the vascular basement membrane – pericytes – mitigate the tumor-supportive nature of the PVN through crosstalk with endothelial cells and macrophages, primarily affecting the glioma molecular phenotype, and secondarily cell proliferation and migration, consequently decelerating tumor progression.

Pericytes have been described to exert pro-tumorigenic functions by contributing to the establishment of immune-tolerance, acting on the ECM composition, and driving epithelial-to-mesenchymal (EMT) transition processes (Hoogstrate et al., 2023; Mader et al., 2018; Valdor et al., 2017; Valdor et al., 2019). Here, we unexpectedly identified pericytes as the most active signaling cell type in both human and murine gliomas. We applied a profoundly characterized mouse model that features the most extensively pericyte-deficient brain microvasculature compatible with adult life observed so far, to study the consequences of pericyte deprivation on glioma development (Andaloussi Mae et al., 2020; Armulik et al., 2010; Nisancioglu et al., 2010). Pericyte depletion led to an aggravated course of tumor development, suggesting a more complex role of pericytes in glioma than described in the literature hitherto. This notion is corroborated by the recent identification of different types of perivascular fibroblasts that might occupy the PVN besides or instead of pericytes (Albiach et al., 2023; Jain et al., 2023; Vanlandewijck et al., 2018) and can have been missed by previous studies. Although the different subtypes of mesenchymal perivascular cells still lack adequate characterization based on thoroughly assorted marker panels, it is conceivable that fibroblasts and pericytes might affect glioma cells and their microenvironments in different ways.

Pathophysiological alterations of the brain vasculature, such as blood-brain barrier dysfunction and neuroinflammation associated with increased immune cell infiltration in the brain, have been observed in neurodegenerative diseases (Cheng et al., 2018; Ozen et al., 2014), SARS-CoV19 infection (Bocci et al., 2021; McQuaid and Montagne, 2023), and demonstrated to be plausibly related to a loss of pericytes in various mouse models featuring a diminished pericyte:endothelial cell ratio (Armulik et al., 2010; Daneman et al., 2010). Furthermore, the generation of subcutaneous tumors in *Pdgfb^ret/ret^*mice led to a more defective tumor vasculature with a hypoxic microenvironment and increased trafficking of MDSCs to the tumor site (Hong et al., 2015). In this study, we did not detect exacerbated hypoxia in pericyte-deprived tumors, but identified altered endothelial cell signaling in the context of an aggravated vascular dysmorphogenesis as a potential driver of glioma progression. One aspect of how the pericyte-poor glioma vasculature fuels tumor growth is the increased number of vessel co-opting tumor cells infiltrating the brain parenchyma. This invasion mechanism, reminiscent of OPC-endothelial migration (Tsai et al., 2016), has been linked to OLIG2^+^/*Wnt7b^+^*tumor cells (Griveau et al., 2018), which is in line with our results indicating OLIG2 positive, OPC-like glioma cells as predominant tumor cells in our model system, and the observed increases in WNT7B signaling of this cell compartment in *Pdgfb^ret/ret^* tumors (Figure S4C). Remarkably, the infiltrative, vessel-associated glioma cells that we observed could potentially acquire mesenchymal features: recently, a study identified a subset of highly vessel co-opting glioma cells, residing midway between the OPC/NPC-like and the mesenchymal cell state on the transcriptomic axis, and linked the induction of this invasive state to angiocrine factors from brain blood vessels (Pichol-Thievend et al., 2024).

In accordance with Hong *et al*., we also measured a higher influx of MDSCs in pericyte-deprived gliomas, which we found paired with an increased attraction of macrophages that had developed an extreme polarization phenotype and could be stratified into subgroups. Spatially, our data showed different preferences of glioma-associated macrophages for the perivascular (*Cd206*^+^ macrophages) and hypoxic (*Cd204^+^* macrophages) niche based on marker gene expression profiles, implying differential activation of tumor-supportive gene programs in the tumor microenvironment; a notion also supported by previous studies (Abdelfattah et al., 2022; Varn et al., 2022; Wang et al., 2024).

The abundance of alternatively activated macrophages has not only been shown to be associated with the MES subtype, but in fact to induce it (Hara et al., 2021) in a manner critical for gliomagenesis and therapy resistance (Abdelfattah et al., 2022; Buonfiglioli and Hambardzumyan, 2021; Sorensen et al., 2018). Here, by integrating scRNA-seq, stRNA-seq, and multiplex imaging data, we identify hypoxic, macrophage-dense tumor zones as the predominant niche for mesenchymal glioma cells (Neftel et al., 2019). We find that whereas most MES tumor cells appear bound to the hypoxic niche, a specific group of extreme MES cells express invasion-related genes, potentially empowering them to leave the hypoxic niche, migrate towards the tumor periphery, and infiltrate the physiologic brain parenchyma. Indeed, in our stRNA-seq data, we observed an increased expression of invasion-related genes at the tumor leading edge. This observation of a high degree of glioma cell infiltration in non-malignant brain parenchyma, emanating from a driving hypoxic/mesenchymal core zone, and advancing through adjacent tiers of proliferating and OPC/NPC/AC like tumor cells, confirms a recent report on the spatial organization of GBM (Greenwald et al., 2024).

Furthermore, we show that MET signaling, a key mesenchymal and cell motility pathway, is activated exclusively in extreme mesenchymal tumor cells of pericyte-poor tumors, and fueled by glioma-associated macrophages that express the ligand HGF, marking a shift in the generally hardwired OPC-like, PDGFB-driven RCAS/tv-a murine glioma model (Ozawa et al., 2014) toward the mesenchymal subtype upon pericyte deprivation. Interestingly, MET activation has already been described for a pericyte-depleted mouse model of breast cancer, and poor pericyte coverage in combination with high *MET* expression has been identified as a predictor of poor outcome in patients with invasive breast cancer (Cooke et al., 2012). However, we do not share the observation of significantly increased hypoxia upon pericyte depletion but surmise instead a remodeled microenvironment as the underlying cause of MET activation. Moreover, AP-1 transcription factor and major regulator of mesenchymal features, FOSL1, a marker of the extreme mesenchymal cells in our dataset, and macrophage secreted HGF, could both potentially induce *Met*/*MET* expression (Cooke et al., 2012; Marques et al., 2021; Pecce et al., 2021; Seol et al., 2000).

The enhancement of mesenchymal features of a subset of GBM cells triggered by alterations of the TME was the most striking alteration upon pericyte depletion in the glioma mouse model that we analyzed. In addition, we also observed subtle changes in tumor composition, such as an expansion of non-tumor oligodendrocyte progenitor cells, a more pronounced NPC-like cell compartment, and an increase of vessel co-opting OLIG2^+^ glioma cells at the tumor leading edge, as well as more dramatic alterations, like the higher abundance of polarized myeloid cells. In this regard, it is interesting to note that longitudinal shifts towards a higher non-tumoral OPC and NPC content and a stronger expression of NPC-like signature genes at the leading-edge, increased tumor cell infiltration and upregulation of immune-suppressive, myeloid-specific expression profiles have been reported for recurring IDH-wildtype gliomas, were found independent of general subtype switches, and could be related to the underlying physical structure and microenvironment of the tumor (Hoogstrate et al., 2023; Varn et al., 2022). Taking also into account that the MES-subtype appears to dominate in recurring gliomas (Hoogstrate et al., 2023; Wang et al., 2022), our data suggest certain parallels regarding the mode of tumor progression between recurrent and pericyte-deprived high-grade gliomas.

In this study, we observed a faster and more aggressive tumor progression in glioma models upon pericyte depletion and subsequent TME remodeling. Our results stress the significance of the TME for the course of glioma progression and its therapeutic potential despite numerous challenges for treatment design due to its complexity. However, they also imply the induction of significant changes in glioma dynamics upon unspecific targeting of pericytes, and consequently demand caution when considering an overall therapeutic ablation of perivascular cells, a concept which is being debated (Cheng et al., 2018; Pombero et al., 2023; Zhou et al., 2017). It is noteworthy that the measured phenotypic shifts towards increased glioma cell invasion and the adoption of the MES cell state are reminiscent of the infiltrative glioma growth pattern that developed upon anti-angiogenic treatment in several studies (Lu et al., 2012; Piao et al., 2013; Piao et al., 2012). To avoid such phenotypic drift, a powerful future integrative treatment approach including the glioma vasculature requires profound knowledge on the diverse subtypes and roles of perivascular stromal cells as a prerequisite to effectively target putative glioma-supporting perivascular cells. At the same time, our results stress the therapeutic importance of preserving pericytes that are contributing to vessel maintenance processes, a notion that is also corroborated by previous findings characterizing pericytes as vulnerable to radiation treatment and linking their ablation to radiation-induced necrosis (Lee et al., 2018), an unfavorable condition that typically induces a mesenchymal switch in glioma cells, making them fit to escape and invade (Falchetti et al., 2019). Finally, it is noteworthy that pericytes are involved in vessel fortification, a beneficial process superior to mere vessel depletion during therapeutic normalization of the glioma vasculature (Martin et al., 2019): the anti-angiogenic drug bevacizumab was shown to increase vessel pericyte coverage and chemotherapy response in breast cancer (Tolaney et al., 2015) and to reduce necrosis in a combination treatment with lomustine (Martin et al., 2019).

Overall, our study reveals the multifaceted nature of pericytes and their role in mitigating high-grade glioma development by orchestrating a tumor-suppressive microenvironment. To achieve improvements in glioma therapy, future studies need to elucidate how specific subgroups of perivascular stromal cells influence the glioma microenvironment and aim to therapeutically target tumor-supportive perivascular cells, while actively shielding pericytes that counteract glioma.

## EXPERIMENTAL PROCEDURES

### KEY RESOURCES TABLE

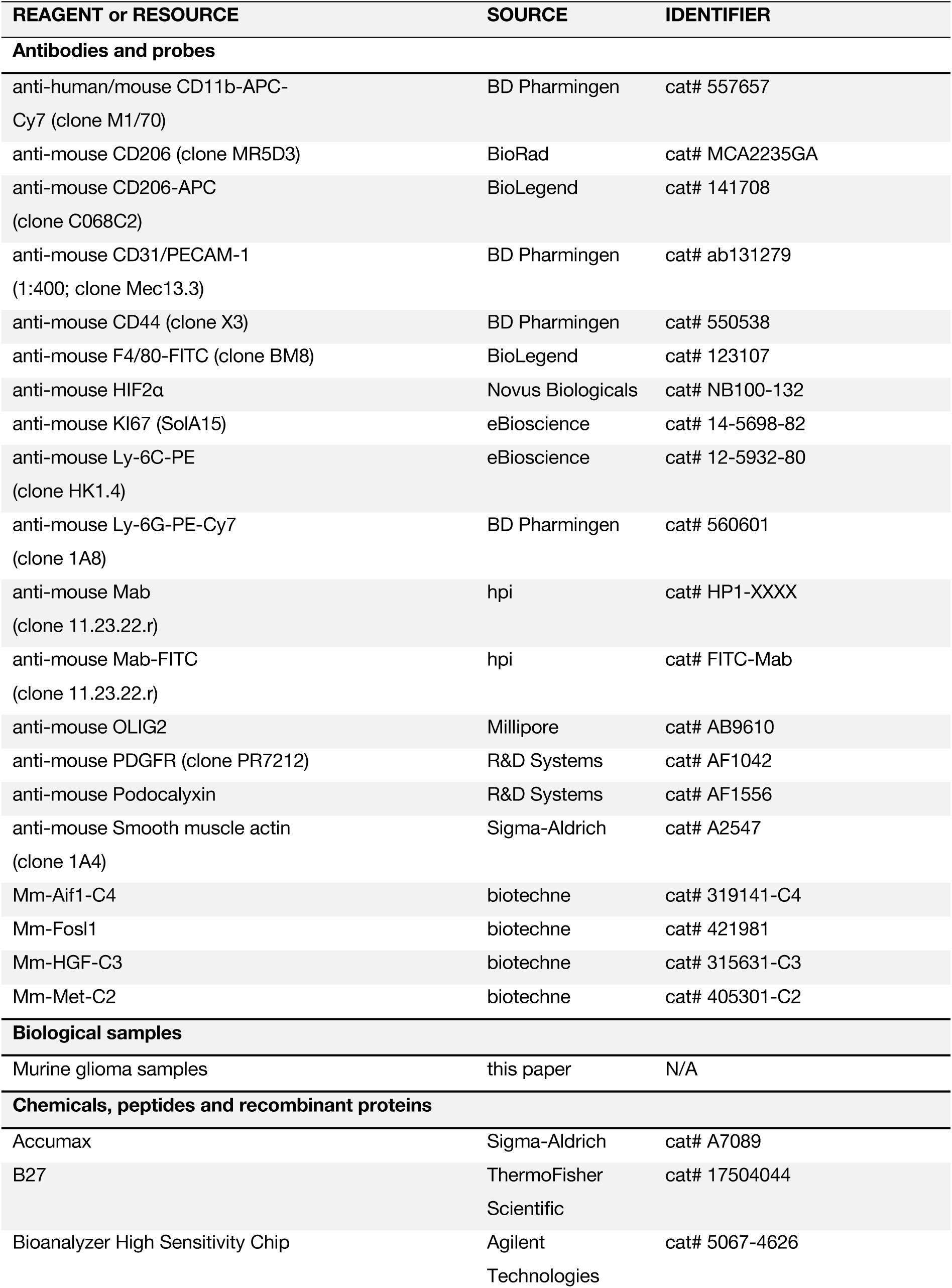
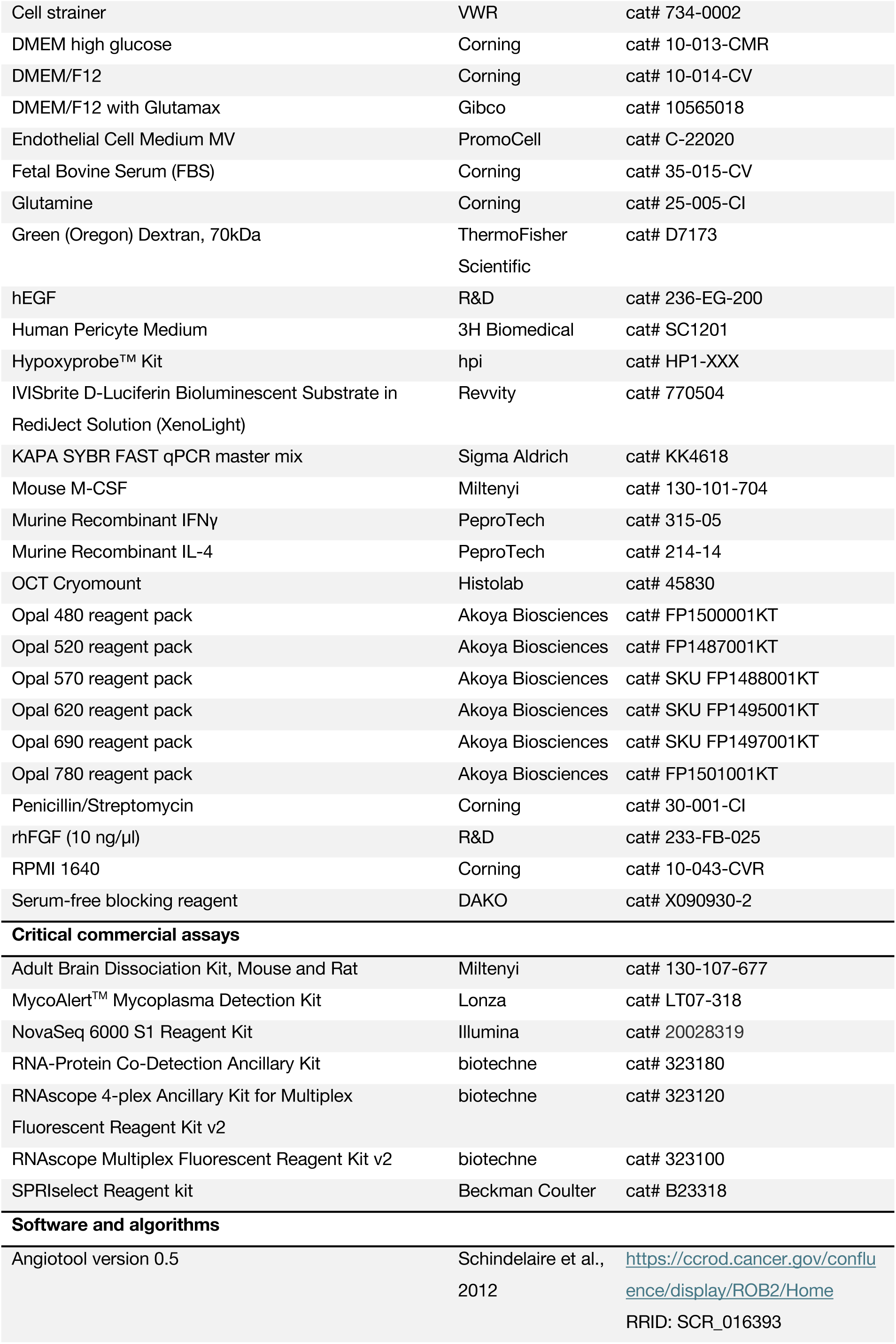
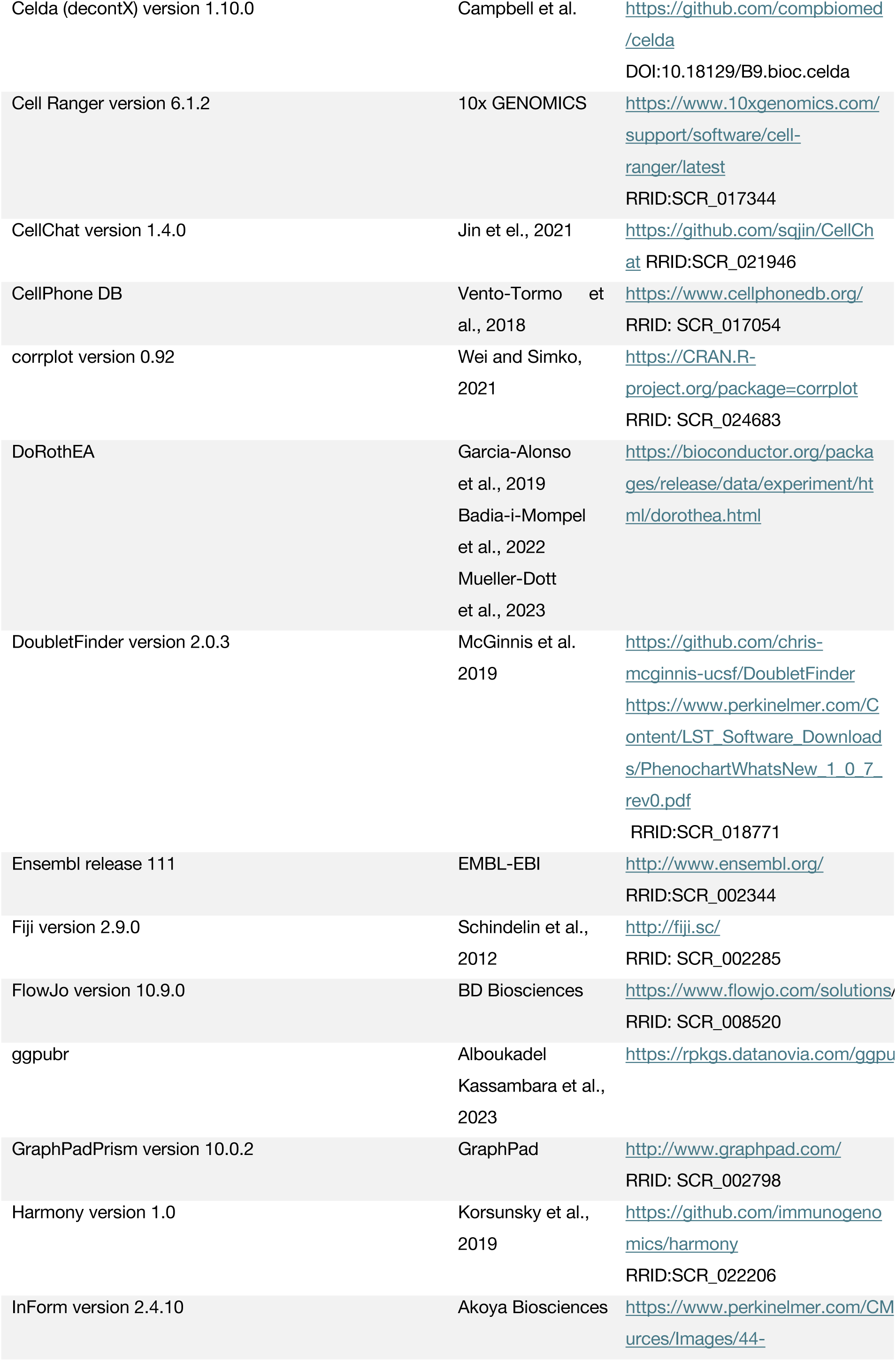
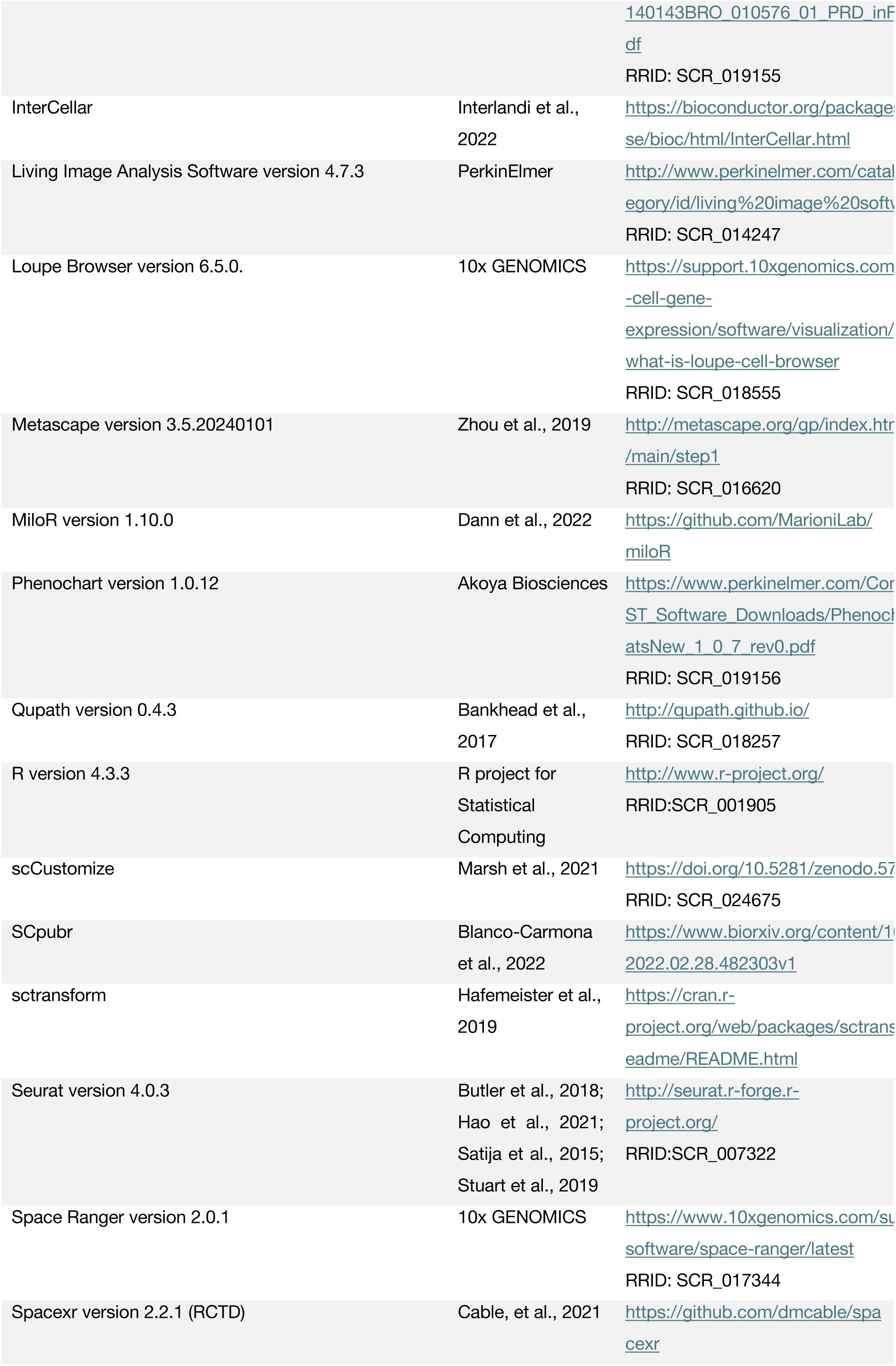
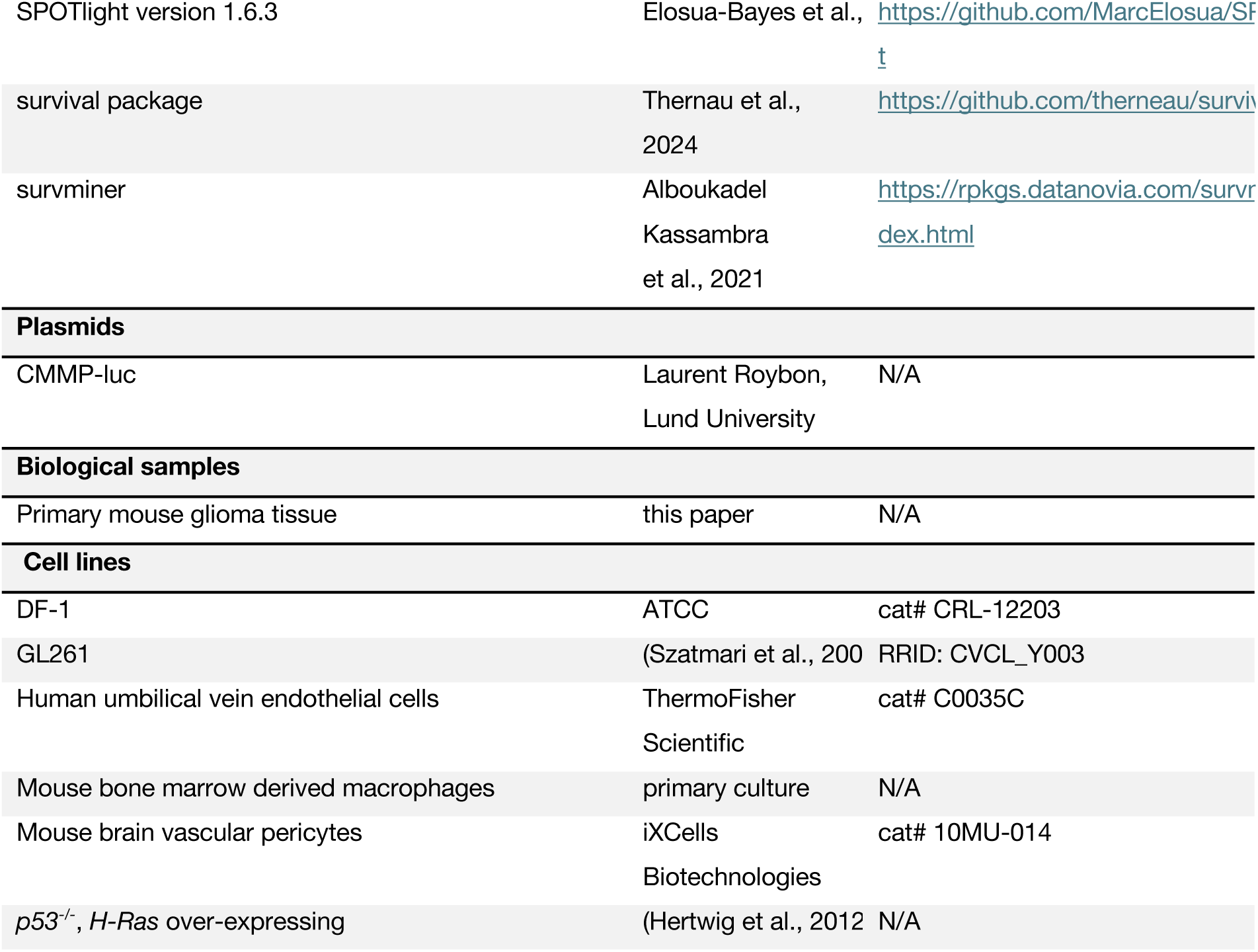

### Generation of murine brain tumors

All animal experiments were performed according to institutional guidelines and approved by the local ethics committee in Lund (permit numbers M167/15, 14122/2020). Transplantations of 100k Gl261, 50k *Tp53^-/-^*, *H-Ras* over-expressing murine glioma cells or 50k MET^+^ or MET^-^ glioma cells into the right frontal brain lope were carried out. For IVIS experiments, 50k *Tp53^-/-^*, *H-Ras* cells expressing a bicistronic GFP-luciferase sequence were transplanted. The RCAS/tv-a system used in this study has been described previously (Holland et al., 1998; Holland and Varmus, 1998; Hu et al., 2005; Uhrbom et al., 2004). *Nestin* Ntv-a, *Pdgfb^ret/+^,* and *Nestin* Ntv-a, *Pdgfb^ret/ret^* mice were used for the RCAS-mediated gliomagenesis in this study. After stereotactical injection of 100k DF-1 virus-producing cells (1:1 mix of RCAS-shp53-and RCAS-PDGFB-transduced cells), mice were observed until they developed brain tumor-related symptoms and euthanized. Mantel-Cox test was applied to test for statistical significance in survival between the different cohorts.

### Immunostaining

For immunostaining, tissues were fresh frozen in OCT Cryomount (Histolab). 10 μm tissue sections were dried at room temperature and fixed in ice-cold acetone for 10 min. All following steps were performed in a humidified chamber. After washing in PBS, sections were blocked with serum-free blocking reagent (DAKO, X090930-2) for 90 min at room temperature. Primary antibodies were diluted 1:100 in PBS + 1% BSA, and sections were incubated overnight at 4 °C. After washing with PBS, secondary antibodies (1:1000 in PBS + 1% BSA) were applied for 1 h at room temperature. Sections were washed and mounted in DAPI-free or DAPI-containing mounting medium. If not stated otherwise, microscopic pictures were taken with an Olympus BX63 microscope.

To determine the pericyte vessel coverage rate, murine glioma sections were immunostained for α-SMA or PDGFRβ respectively, and PODXL. 22 fields of view (FOV) of three *Pdgfb^ret/ret^*and four *Pdgfb^ret/+^* animals were analyzed for Gl261-derived gliomas and 200 FOV of three *Pdgfb^ret/ret^* and five *Pdgfb^ret/+^* animals were analyzed for PDGFB-induced gliomas. Pericyte coverage was calculated by dividing the total α-SMA or PDGFRβ positive area by the total PODXL positive area, applying Angiotool (version 0.5) (Zudaire et al., 2011) and Fiji (version 2.9.0) (Schindelin et al., 2012).

### Characterization of the murine glioma tissue

To assess vessel parameters, glioma sections of 7 *Pdgfb^ret/ret^*and 7 *Pdgfb^ret/+^* mice were immunostained for CD31, and 9-10 FOV per mouse were analyzed with Fiji.

For vessel perfusion analysis, mice were briefly sedated with isoflurane and injected with 40 μl Oregon-Green Dextran (70 kDa, 25 mg/ml, ThermoFisher Scientific) in a retro-orbital fashion. After 15 min incubation, animals were subjected to cardiac perfusion with 10 ml PBS and 10 ml 4% pFA, and brains were collected subsequently.

For hypoxia and proliferation analysis, glioma sections were immunostained against HIF-2α and KI67, counterstained with DAPI and the HIF-2α/DAPI and KI67/DAPI total area ratios determined.

For hypoxia analysis with the Hypoxyprobe system (hpi), animals were injected in a retro-orbital fashion with 50 μl pimonidazole (60 mg/kg body weight), euthanized after 30 min incubation, and brains collected.

To analyze the invasive rim (IR) zone, glioma cryosections of 3 *Pdgfb^ret/ret^* and 5 *Pdgfb^ret/+^* mice were immunostained for OLIG2 and PECAM, and evaluated subsequently. 5 FOV/section were analyzed by assessing the border of the tumor bulk and counting of OLIG2^+^ cells, PECAM1^+^ cells, OLIG2^+^ positive cells in direct contact with PECAM1^+^ cells, as well as determining the total PECAM1^+^ area in the adjacent IR zone with Fiji.

### *In vivo* bioluminescence imaging of glioma development in mice

Murine *p53^-/-^*, *H-Ras* over-expressing glioma cells were transduced with a CMMP vector (provided by Laurent Roybon, Lund University, Lund, Sweden) containing the luciferase cDNA sequence. Three-to six-months-old *Pdgfb^ret/ret^* and *Pdgfb^ret/+^* mice were subjected to stereotactical engraftment of 100k luciferase over-expressing glioma cells into the right frontal brain lope. From five days after engraftment, the tumor growth was monitored by non-invasive 2D bioluminescence (BLI) imaging, using IVIS-CT spectrum (PerkinElmer). Briefly, mice were anesthetized with 3% isoflurane gas and injected intraperitoneally with 150 mg D-Luciferin/kg of body weight (Revvity) in PBS prior to imaging. Acquisition of 2D images was performed sequentially with a five-minute interval between different segments of exposure (emission: open filter, f/stop: 1, binning: 8). BLI signal intensity was quantified in total flux (photons/s) after deducting the average background signal (Bkg) from measurement regions of interest (ROI), using the Living image analysis software (version 4.7.3, PerkinElmer). To estimate tumor growth kinetics, the data was fitted to a non-linear model (least squares fit) using GraphPadPrism (version 9.0, GraphPadSoftware).

### Cell culture

All cells were cultured at 37 °C and 5% CO_2_, and were frequently checked for mycoplasma infections using the MycoAlert^TM^ Mycoplasma Detection Kit (Lonza). Unless stated otherwise, all media were supplemented with glutamine (2 mM, Corning), Penicillin/Streptomycin (50 μg/ml, Corning), HEPES (25 mM, Corning) and 10% FBS (Corning). DF1 cells were cultured in DMEM high glucose (Corning). Macrophages were cultured in RPMI 1640 glucose (Corning) supplemented with 50 ng/ml M-CSF (Miltenyi). Glioma cells were cultured under serum-free conditions as spheres, in medium supplemented with B27 (ThermoFisher Scientific, 17504044), rhFGF (10 ng/μl, R&D, 233-FB-025) and hEGF (10 ng/μl, R&D, 236-EG-200) in DMEM/F12 (Gibco). Pericytes were cultured in pericyte medium containing relevant growth factors (3H Biomedical, 1201).

### TSP assay

2000 cells per 2 ml medium were seeded in 6-wells in triplicates. One week after incubation, spheres were counted with a Zeiss Axio Vert.A1 system. Subsequently, spheres were digested with Accumax (Sigma-Aldrich), triturated, and cells counted with a Buerker chamber. 2000 cells per triplicate were seeded again for continuous analysis.

### Macrophage assay with pericyte-conditioned medium

Pericytes and endothelial cells at 80% confluency were cultured in macrophage medium w/o growth factors for 48 h. Subsequently, the medium was collected, centrifuged, filtered, and applied to macrophages at 50-80% confluency in 6-wells. M-CSF (Miltenyi) was added to all wells and IFNγ (14 ng/ml, PeproTech) or IL-4 (14 ng/ml, PeproTech) was added to the respective sample wells. After medium exchange, the macrophages were cultured for 48 h, scraped off and frozen at -150°C in cryopreservation medium (70% medium, 20% FBS, 10% DMSO).

### Glioma sample collection

Terminally ill animals were sedated with isoflurane, decapitated, and the brains removed. Brains were immediately brought on ice, and a coronal cut through the tumor lesion was performed. Tumor material was then isolated and washed in ice-cold DPBS. Subsequently, the tumor tissue was dissociated using the Adult Brain Dissociation Kit (Miltenyi), resuspended in cryopreservation medium, and stored at -150°C.

### FACS

Cells were thawed quickly in a 37°C warm water bath, resuspended in medium, and centrifuged at 220 g. Subsequently, cells were washed twice with 10 ml FACS buffer (DPBS with 5% FBS or BSA), centrifuged at 220 g and resuspended in 50-100 μl FACS buffer at a concentration of 1:100 per respective antibody. After 30 min incubation on ice in the dark, the cells were washed again in FACS buffer, passed through a 40 μm cell strainer (VWR), spun at 220 g, and resuspended in FACS buffer at a density of 1-5x10^6^ per 0,5 ml. The cells were then placed on ice and kept in the dark until analysis with a BD FACS Melody system (BD Biosciences). Post-FACS analysis of the recorded data was performed with FLOWJO (version 10.9.0, BD Biosciences). Sorted cells were centrifuged at 300 g, resuspended in 500 μl growth medium, and initially cultured in 24 wells.

### Single-cell RNA sample preparation and sequencing

High-grade gliomas were initiated by stereotactically delivering the glioma oncogenic drivers PDGFB and shp53 into the subventricular zone of transgenic adult *Ntv-a* mice that were either pericyte-poor (*Pdgfb^ret/ret^*) and or control (*Pdgfb^ret/+^*). 7 *Pdgfb^ret/+^* and 4 *Pdgfb^ret/ret^* mice were sacrificed when displaying severe symptoms. Subsequently, whole tumors were extracted and processed into single cell suspensions according to the protocol of the Adult Brain Dissociation Kit, mouse and rat, (Miltenyi). In short, whole tumor was minced into smaller slices with a scalpel and incubated for 30 min with the supplier’s enzyme mixes, while being heated at 37°C and mechanically grinded in the gentle MACS Octo Dissociater with Heaters (Miltenyi). Subsequently, the sample was resuspended and applied to a 70 μm MACS SmartStrainer (Miltenyi), after which the single cell suspension underwent subsequent steps for debris removal and lysis of erythrocytes. For library preparation, 18,000 cells / 45 μl in 0.04% BSA in PBS were delivered to the Center of Translational Genomics at Lund University (CTG). Single-cell 3’ RNA-seq libraries were prepared using 10x Chromium system (v3.1) by CTG. Samples were run using Read 1 28 cycles, i7 index 5 cycles, i5 index 0 cycles, Read 2 91 cycles on the Illumina NovaSeq 6000 to a depth of 50,000 reads/cell. This was performed in two different batches (time points). The first batch included 2 *Pdgfb^ret/+^* and 1 *Pdgfb^ret/ret^* mice, while the second batch included 5 *Pdgfb^ret/+^* and 3 *Pdgfb^ret/ret^*mice.

### Single-cell RNA data analyses

The raw reads were processed, mapped, and quantified using Cell Ranger count (v6.1.2) (Zheng et al., 2017) with default settings, with an initial expected cell count of 10,000. We mapped the reads to a modified mouse reference genome (mm10) that contained the RCAS-PDGFB longest possible sequence following Cell Ranger instructions. Cell Ranger’s filtered files were further filtered to keep cells with number of transcripts lower than 50,000, number of features larger than 250 and lower than 8000, and percentage of mitochondrial and ribosomal genes lower than 10% and 25%, respectively. Finally, we filtered out mitochondrial and hemoglobin genes, as well as features with less than 20 counts across cells. We used DoubletFinder (v2.0.3) to filter doublets assuming a doublet formation rate of 7.5%. pN=0.25 and pK were adjusted for each sample using pN-pK parameter sweeps and mean-variance-normalized bimodality coefficient (BCmvn) using paramSweep_v3 (McGinnis et al., 2019). We used DecontX (celda, v1.6.1) to filter ambience DNA contaminations (Yang et al., 2020). We filtered out barcodes that had more than 25% of reads identified as ambient RNA contamination. QC and filtering steps were performed in every sample separately. Data analyses, including normalization, scaling, dimensionality reduction (PCA) and clustering was performed using Seurat (v4.0.3) (Butler et al., 2018; Hao et al., 2021; Satija et al., 2015; Stuart et al., 2019). Batch integration was performed using the Harmony algorithm v1.0, specifying batch and sample (Korsunsky et al., 2019). The Seurat function FindConservedMarkers() was used for finding differentially expressed genes between clusters, and for testing for differential expression between conditions for every cluster separately. The top 50 DEG for every cluster were defined as the cluster/cell type signature (Table S1). We used scCustomize (v2.1.2) (Marsh SE (2021). scCustomize: Custom Visualizations & Functions for Streamlined Analyses of Single Cell Sequencing) for visualization of the results, including UMAP, feature plots and clustered dot plots of selected top differentially expressed genes. scCustomize uses k-means method for clustering dot plots. For the butterfly plot visualization, the AddModuleScore function in Seurat was used to calculate the Neftel *et al*. gene set enrichment/module scores (AC-like, MES-like, NPC-like, and OPC-like as in the original publication) and the do_CellularStatesPlot function in SCpubr (v2.0.2) (Blanco-Carmona, E., Generating publication ready visualizations for Single Cell transcriptomics using SCpubr. bioRxiv, 2022: p. 2022.02.28.48230) was used to construct the scatter plots. Transcription factor activity analysis was performed using DoRothEA (v1.6.0) (Badia et al., 2022; Garcia-Alonso et al., 2019). Transcription factor regulons with the three highest confidence levels (“A”, “B”, and “C”) were used and the averaged activity scores per cluster were plotted using the do_TFActivityPlot function in the SCpubr package. We used default parameters for these tools unless otherwise specified.

### Z-score estimation and correlation matrix

To further explore gene expression patterns in our dataset and compare our defined gene signatures to human defined counterparts, we used published available gene signatures from Neftel et al. and Couturier et al. Briefly, we used Neftel et al. gene signatures as reported by the authors: AC, OPC, MES1, MES2, NPC1, NPC2, G1S, G2M. The gene lists reported by Couturier et al. are from a continuous scale. Therefore, we selected the top 20 and 20 bottom genes of their eigenvectors DC1, DC2 and DC3, and defined 6 Couturier signatures as DC1 top non-neuronal-like (DC1t/non-neuronal), DC2 top oligodendrocyte (DC2t/OLC), DC3 top progenitor (DC3t/progenitor), DC1 bottom neuronal-like (DC1b/neuronal), DC2 bottom Mesenchymal-Astrocyte-like (DC2b/Mes-Astro) and DC3 bottom non-progenitor (DC3b/non-progenitor). From these genes, we identified 1:1 orthologs between mouse and human using an orthology table downloaded from Ensemble (release 111) using the Biomart web-based tool. The exact genes included are reported in Table S2. For every gene signature, calculation of Z-scores was performed for each gene belonging to the signature. Z-scores enabled us to estimate how many standard deviations from the mean a value is. Considering that x_ij_ is the expected value of gene i in sample j, μi is the mean of expected values for gene i across all j samples and σ_i_ is the standard deviation of expected values for gene i in all j samples, the formula of the Z-score is:

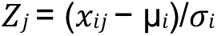

Signature estimation value for every cell is then performed by calculating the arithmetic mean of Z-scores of all the genes included in the signature. We estimated pairwise Pearson’s correlation coefficients for every pair of signatures across all cells. A correlation matrix was calculated and plotted using the corrplot R package (v0.92) (Wei T, Simko V, 2021). The correlation matrix plot was ordered using hierarchical clustering for better visualization. Black squares were manually added to highlight selected signature clusters.

### Differential cell abundance

Changes in cell type abundance between *Pdgfb^ret/+^ and Pdgfb^ret/ret^* derived cells was calculated for every cluster as:

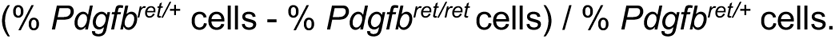

We also used MiloR (v1.1) (Dann et al., 2022) to estimate changes in cell type abundance. Instead of using discrete clusters, MiloR tests for differential abundance in neighborhoods derived from a k-nearest neighbor graph. The parameters used were prop = 0.05, k = 30, d = 20, alpha=0.2.

### Human glioma single-cell RNA data and spatial transcriptomics analyses

The human single cell RNA-seq dataset (Abdelfattah et al., 2022) comprising 18 glioma samples was downloaded from Broad Institute’s Single Cell Portal (https://singlecell.broadinstitute.org/single_cell). The dataset was then processed using the basic pipeline of Seurat. The cell-type annotations and UMAP coordinates were provided by the authors. Clusters and results from gene expression level analyses were visualized using the scCustomize package.

The previously published human GBM spatial transcriptomic data set (Ravi et al., 2022) was downloaded from https://datadryad.org/stash/dataset/doi:10.5061/dryad.h70rxwdmj. Data (tumor #UKF 269_T) normalization using SCTransform, principal component analysis and further processing were all done in Seurat using default parameters. Seurat’s AddModuleScore function was used to calculate gene set module scores. The module scores were visualized onto the Visium tissue section using Seurat’s SpatialFeaturePlot function. The module scores were also aggregated by the different histological areas of the sample and visualized in heatmaps using the do_EnrichmentHeatmap function as provided in the SCpubr package.

### Cell-cell communication analyses

Cell-cell communication strength is modeled as the probability of ligand-receptor interaction based on the scRNA-seq gene expression data of cells, in conjunction with a database containing prior knowledge of ligands, receptors and cofactors. To analyze the communication routes between the cell populations in the human glioma dataset, CellphoneDB (v2.0.0) (Efremova et al., 2020) was used to characterize the number of interactions and CellChat (v1.4.0) (Jin et al., 2021) was used to calculate the interaction number and strength using default parameters. Results were visualized using CellChat, ggpubr (v0.6.0) and InterCellar (v2.0.0). To identify the conserved and context-specific signaling between *Pdgfb^ret/+^* and *Pdgfb^ret/ret^* pathways, we performed a comparative analysis using CellChat (Jin et al., 2021) . Cell-cell communication analyses were performed separately for *Pdgfb^ret/+^* and *Pdgfb^+/+^* derived cells, and then compared.

### Spatial transcriptomics sample preparation

Sample preparation was done according to the protocol "Tissue preparation guide" (CG000240; 10x GENOMICS). All steps were performed using the 10x Visium Spatial Gene Expression Slide & Reagent kit. Methanol fixation and immunofluorescence staining were done according to protocol CG000312. The optimal tissue optimization time (4 min) was determined in a time course experiment according to protocol CG000238RevE. cDNA synthesis and library generation were done according to protocol CG000239RevD. cDNA synthesis was carried out according to the results obtained from qPCR testing with a KAPA FAST SYBR qPCR system (KAPA Biosystems). The amplified cDNA was processed according to protocol, using the SPRIselect Reagent kit (Beckman Coulter). Quality control of the final libraries was performed on an Agilent Bioanalyzer using the High Sensitivity DNA assay (Agilent Biotechnologies). Libraries were sequenced on NovaSeq 6000 (Illumina) using the S1 100 reagent kit (Illumina) with 1% PhiX, and the following sequencing parameters: Read 1 – 28 cycles, Read 2 – 90 cycles, Index 1 – 10 cycles, Index 2 – 10 cycles.

### Image pre-processing and sequencing analyses

Visium expression slides were imaged on a Leica DMi8 microscope. Images were recorded using the Leica Application Suite (LAS X) software, and computationally cleared with the THUNDER technology (Leica). Images were then imported into -, where brightness and contrast were adjusted separately for every channel. Adjusted images were saved as a single multi-stack composite .TIFF file for every sample separately. These .TIFF images were imported into the 10x Genomics Loupe Browser 6.5.0 for manual alignment. We also saved a separate .TIFF file containing channel 3 (PE/PIM), where brightness was adjusted until the fiducial frames (FF) became visible. These images were imported into the Loupe browser as an auxiliary for FF alignment. Manual alignment of the FF, and manual tissue selection was performed in 10x Genomics Loupe Browser 6.5.0, following 10X instructions and recommendations (online Manual Fiducial Alignment guide for Visium). We used Space Ranger count (v2.0.1) to map and count sequencing reads, using the modified reference genome (mm10) containing the RCAS-PDGF sequence (created for mapping the scRNA-seq reads). The multi-stack composite TIFFs (using the darkimage option) and JSON alignments files were used as input for Space Ranger count.

### Spatial transcriptomics data analysis

We used the filtered feature matrix from Space Ranger as an input for downstream analysis. In short, we removed any spots with 0 counts, fewer than 500 genes detected, mitochondrial content higher than 25%, and percent hemoglobin genes detected higher than 20%. We also filtered out mitochondrial and hemoglobin genes. QC and filtering steps were performed in every sample separately. Data analyses, including normalization (logNormalize), clustering and differential gene expression was performed using Seurat toolkit (v4.0.3) (Butler et al., 2018; Hao et al., 2021; Satija et al., 2015; Stuart et al., 2019). Batch integration was performed using the Harmony algorithm (v1.0) (Korsunsky et al., 2019), specifying batch and sample-id as covariates. The function FindConservedMarkers() was used for finding differentially expressed genes between clusters. We used the Robust Cell Type Decomposition (RCTD) (Cable et al., 2022) method, a maximum-likelihood approach, with ‘full mode’ for estimating cell type proportions on spatial transcriptomics spots. We used our scRNA-seq dataset as a reference data set to deconvolve every spot in our 8 spatial transcriptomics samples. The RCTD output is a weighted proportion (0-1), meaning that the sum of all the cell type proportions equals to 1 in every spot. We used the RCTD proportions and the coordinates of each spot as input to the function plotSpatialScatterpie() from the SPOTlight package (v 1.6.3) (Elosua-Bayes et al., 2021) to create a spatial scatterpie chart.

### Patient cohort analyses

TCGA GBM gene expression profiles and matched clinical information were retrieved from the cBioPortal database (https://pubmed.ncbi.nlm.nih.gov/22588877/) and https://pubmed.ncbi.nlm.nih.gov/23550210/. The TCGA RNA-seq dataset was used for the expression analysis of the primary and recurrent GBM samples, and for the survival analysis, the TCGA Affymetrix HT HG U133A expression dataset was used. Overall survival (OS) and progression-free interval (PFI) data were used to analyze clinical outcomes. The survival analysis was performed using the Surv and survfit functions in the survival package (v3.5-5) and the ggsurvplot function in survminer (v0.4.9). We stratified the patients into two expression groups (low and high) according to the median value of the mean expression of the gene signatures. Kaplan-Meier survival curves and log-rank tests were used to evaluate the performance of the gene signatures.

## Acknowledgements

KP is the Grosskopf Professor of Molecular Medicine at Lund University. This work was supported by grants from the Swedish Research Council, the Swedish Cancer Society, the Swedish Childhood Cancer Society, the Knut and Alice Wallenberg foundation, Swedish State Support for Clinical Research through Region Skåne ALF, the Göran Gustafsson foundation, the Mats Paulsson foundations, the Cancera foundation.

We thank the Centre for Translational Genomics (Lund University), and Clinical Genomics Lund (SciLifeLab) for sequencing services; Uppsala Multidisciplinary Center for Advanced Computational Sciences (UPPMAX), and the Swedish National Infrastructure for Computing (SNIC) for computing resources. The authors would like to thank Ulrike Nuber for kindly providing the *p53^-/-^*, *H-Ras* over-expressing glioma cell line, Laurent Roybon for kindly providing the CMMP vector, Massimo Squatrito for kindly providing the Ntv-a mice and Christer Betsholtz as well as Guillem Genové for kindly providing the *Pdgfb^ret/ret^* mice. The authors would like to thank Wondossen Sime, David Lindgren, Eliane Cortez and Christina Möller for helpful technical advice and assistance.

## Author Contributions

Conceptualization: KP, SB, AP. Methodology: PB, CO, JS, MB, KH, MST, BP, EC, RR, EJ, GBJ, SB. Software: PB, JS. Validation: CO, JS, PB, SB. Formal analysis: PB, CO, JS, KH, KP, SB. Investigation: PB, CO, JS, KH, BP, EC, SB. Resources: GBJ, AP, KP. Data curation: PB, CO, JS, SB, KP. Writing – original draft: SB, KP, PB, JS. Writing – review & editing: PB, CO, JS, MB, KH, MST, BP, EC, RR, EJ, GBJ, AP, KP, SB. Visualization: PB, JS, CO, SB. Supervision: KP, SB. Project administration: KP, SB. Funding acquisition: KP.

