## Supplemental materials for "Pericytes orchestrate a tumor-restraining microenvironment in glioblastoma"

### Figure S1

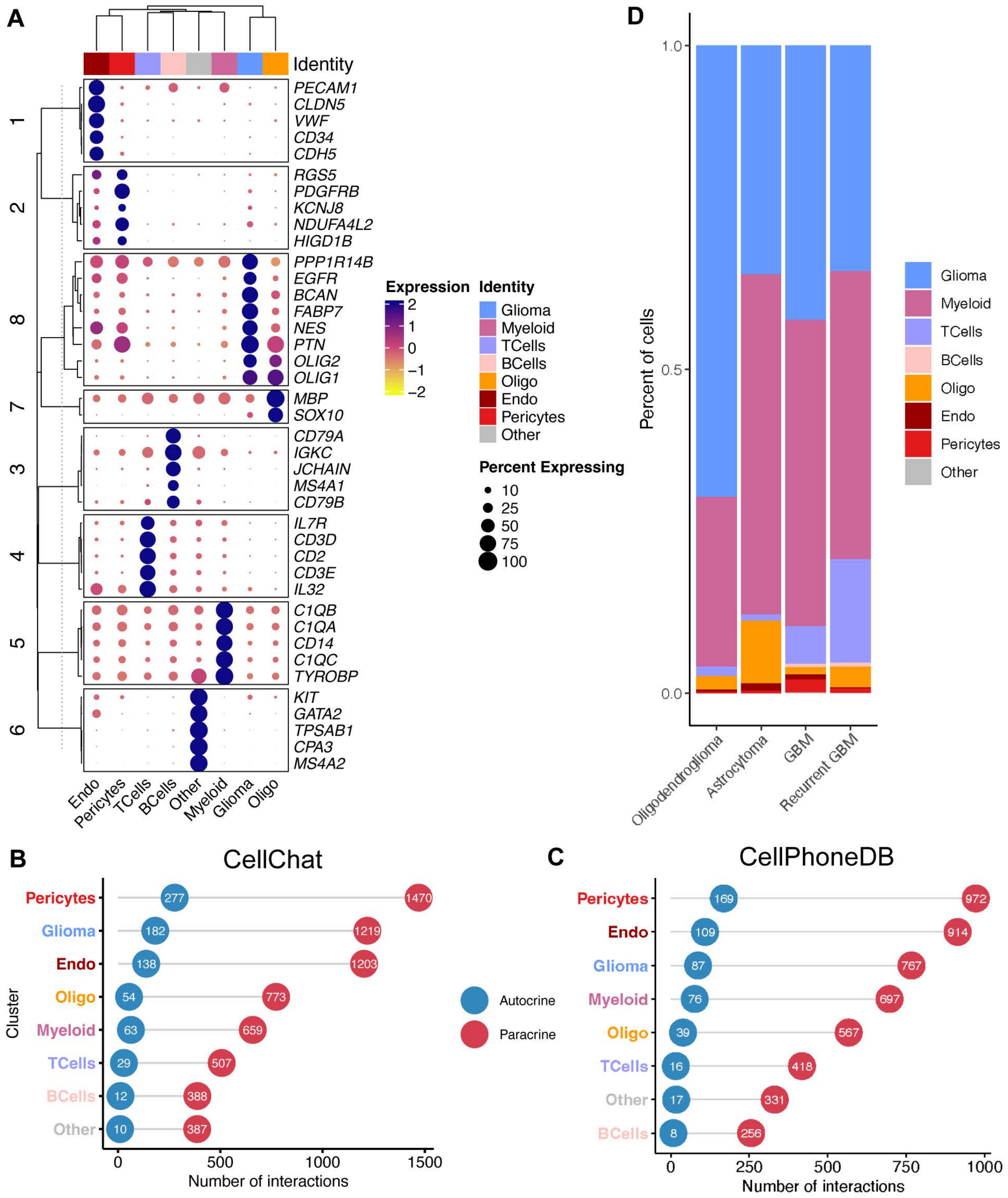

A
Identity
1
2
8
7
3
4
5
6
PECAM1
CLDN5
VWF
CD34
CDH5
RGS5
PDGFRB
KCNJ8
NDUFA4L2
HIGD1B
PPP1R14B
EGFR
BCAN
FABP7
NES
PTN
OLIG2
OLIG1
MBP
SOX10
CD79A
IGKC
JCHAIN
MS4A1
CD79B
IL7R
CD3D
CD2
CD3E
IL32
C1QB
C1QA
CD14
C1QC
TYROBP
KIT
GATA2
TPSAB1
CPA3
MS4A2
Endo
Pericytes
TCells
BCells
Other
Myeloid
Glioma
Oligo
Expression
2
1
0
−1
−2
Identity
Glioma
Myeloid
TCells
BCells
Oligo
Endo
Pericytes
Other
Percent Expressing
10
25
50
75
100
D
Percent of cells
1.0
0.5
0.0
Oligodendroglioma
Astrocytoma
GBM
Recurrent GBM
Glioma
Myeloid
TCells
BCells
Oligo
Endo
Pericytes
Other
B
CellChat
Cluster
Pericytes
Glioma
Endo
Oligo
Myeloid
TCells
BCells
Other
277
1470
182
1219
138
1203
54
773
63
659
29
507
12
388
10
387
0
500
1000
1500
Number of interactions
Autocrine
Paracrine
C
CellPhoneDB
Pericytes
Endo
Glioma
Myeloid
Oligo
TCells
Other
BCells
169
972
109
914
87
767
76
697
39
567
16
418
17
331
8
256
0
250
500
750
1000
Number of interactions

**Figure S1. Characterization of different cell groups isolated from human glioma samples, related to Figure 1.**

(A) Clustered dot plot showing the scaled average expression of selected top differentially expressed genes across all clusters identified in the scRNA-seq dataset from Abdelfattah et al.
(B-C) Lollipop plots showing the number of autocrine and paracrine interactions associated with each cell population for human glioma samples of the scRNA-seq dataset from Abdelfattah et al., generated with CellChat (B) and CellPhoneDB (C).
(D) Bar plot indicating the relative abundance of annotated cell groups in human glioma samples (dataset of Abdelfattah et al.).

### Figure S2

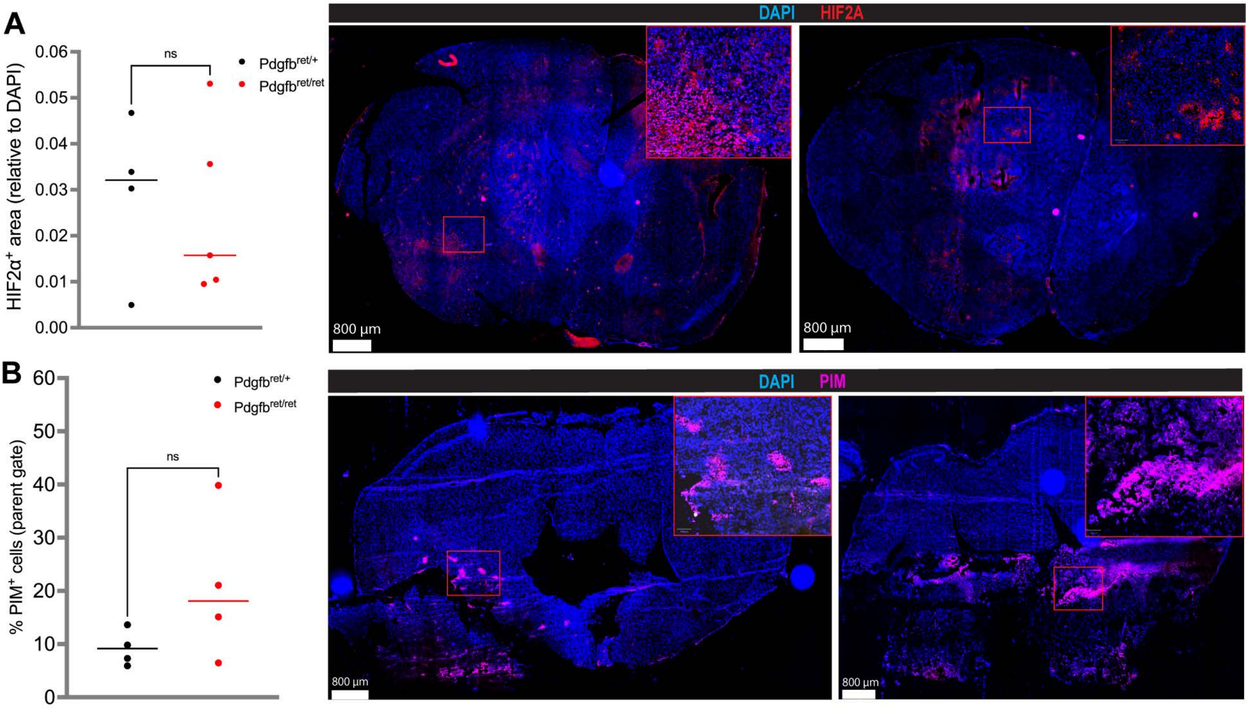
A
B
HIF2α+ area (relative to DAPI)
0.06
0.05
0.04
0.03
0.02
0.01
0.00
ns
Pdgfbret/+
Pdgfbret/ret
DAPI
HIF2A
800 µm
800 µm
% PIM+ cells (parent gate)
60
50
40
30
20
10
0
ns
Pdgfbret/+
Pdgfbret/ret
DAPI
PIM
800 µm
800 µm

**Figure S2. Hypoxia quantification comparing *Pdgfb*$^{ret/+}$ and *Pdgfb*$^{ret/ret}$ murine glioma, related to Figure 2.**

(A) Hypoxia analysis (left panel), based on the quantification of HIF2α immunostainings (middle and right panel).
(B) Hypoxia analysis, based on the quantification of PIM$^+$ cells with FACS (left panel). The middle and right panel show representative immunostainings of PIM.
Boxes denote the enlarged regions. ns not significant

### Figure S3

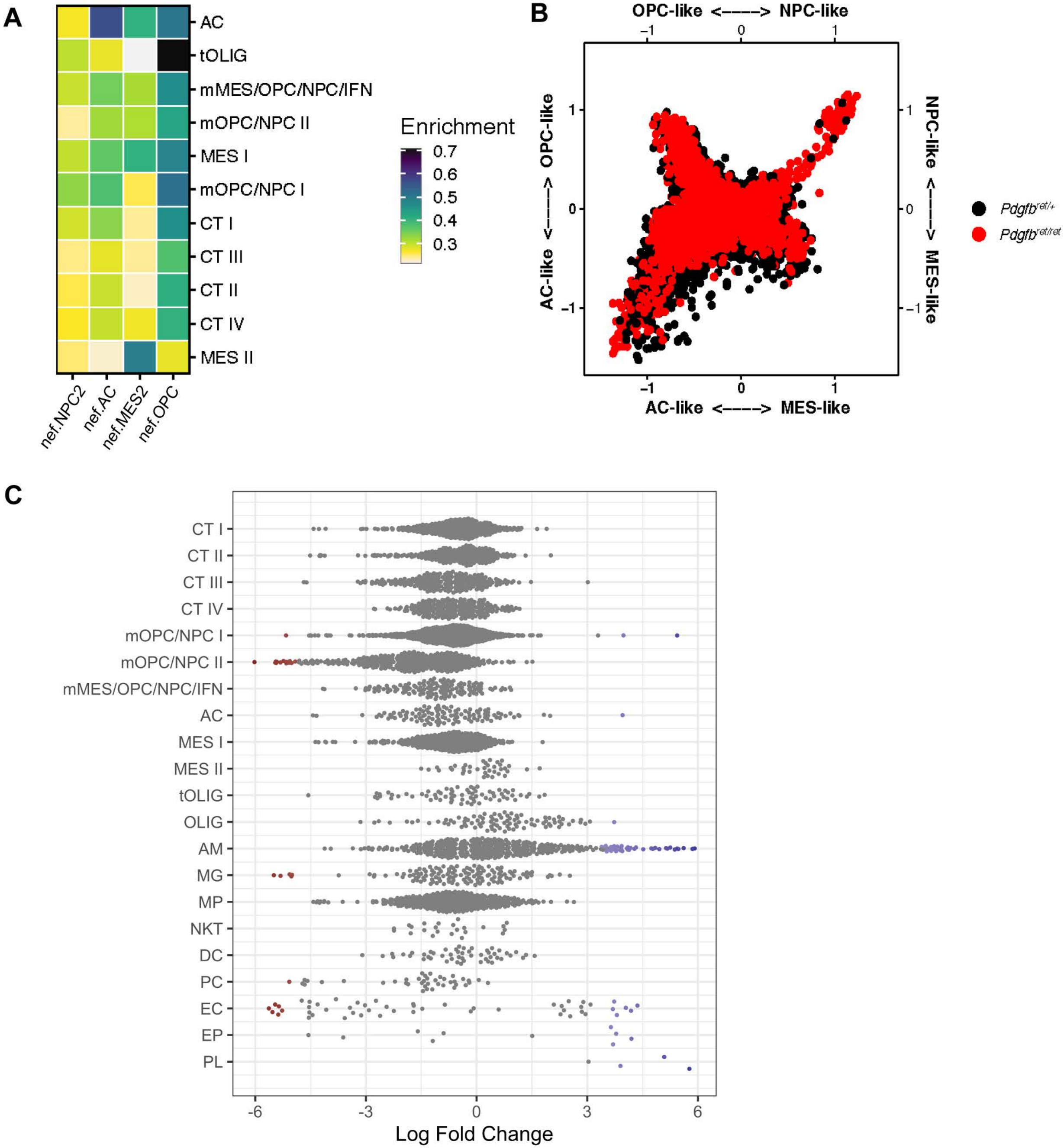

A
AC
tOLIG
mMES/OPC/NPC/IFN
mOPC/NPC II
MES I
mOPC/NPC I
CT I
CT III
CT II
CT IV
MES II
nef.NPC2
nef.AC
nef.MES2
nef.OPC
Enrichment
0.7
0.6
0.5
0.4
0.3
B
OPC-like <----> NPC-like
AC-like <----> OPC-like
NPC-like <----> MES-like
AC-like <----> MES-like
Pdgfb^ret/+
Pdgfb^ret/ret
C
CT I
CT II
CT III
CT IV
mOPC/NPC I
mOPC/NPC II
mMES/OPC/NPC/IFN
AC
MES I
MES II
tOLIG
OLIG
AM
MG
MP
NKT
DC
PC
EC
EP
PL
-6
-3
0
3
6
Log Fold Change

**Figure S3. Abundance and distribution of cell populations in RCAS-induced glioma, related to Figure 3.**

(A) Heatmap representing the enrichment of the molecular subtype NPC-like, OPC-like, AC-like and MES-like signatures from Neftel *et al.* in the murine glioma scRNA-seq dataset.

(B) Butterfly plot of the molecular subtype signature scores from Neftel *et al.*, applied to the murine glioma scRNA-seq data, and comparing the relative signature scores of all $\textit{Pdgfb}^{ret/+}$ and $\textit{Pdgfb}^{ret/ret}$ glioma sample cells.

(C) Results from MiloR differential abundance test. Beeswarm plot of the distribution of log fold changes between $\textit{Pdgfb}^{ret/+}$ and $\textit{Pdgfb}^{ret/ret}$ derived neighborhoods from different cell types. Differential abundance neighborhoods at FDR 20% are colored.

**Figure S4**

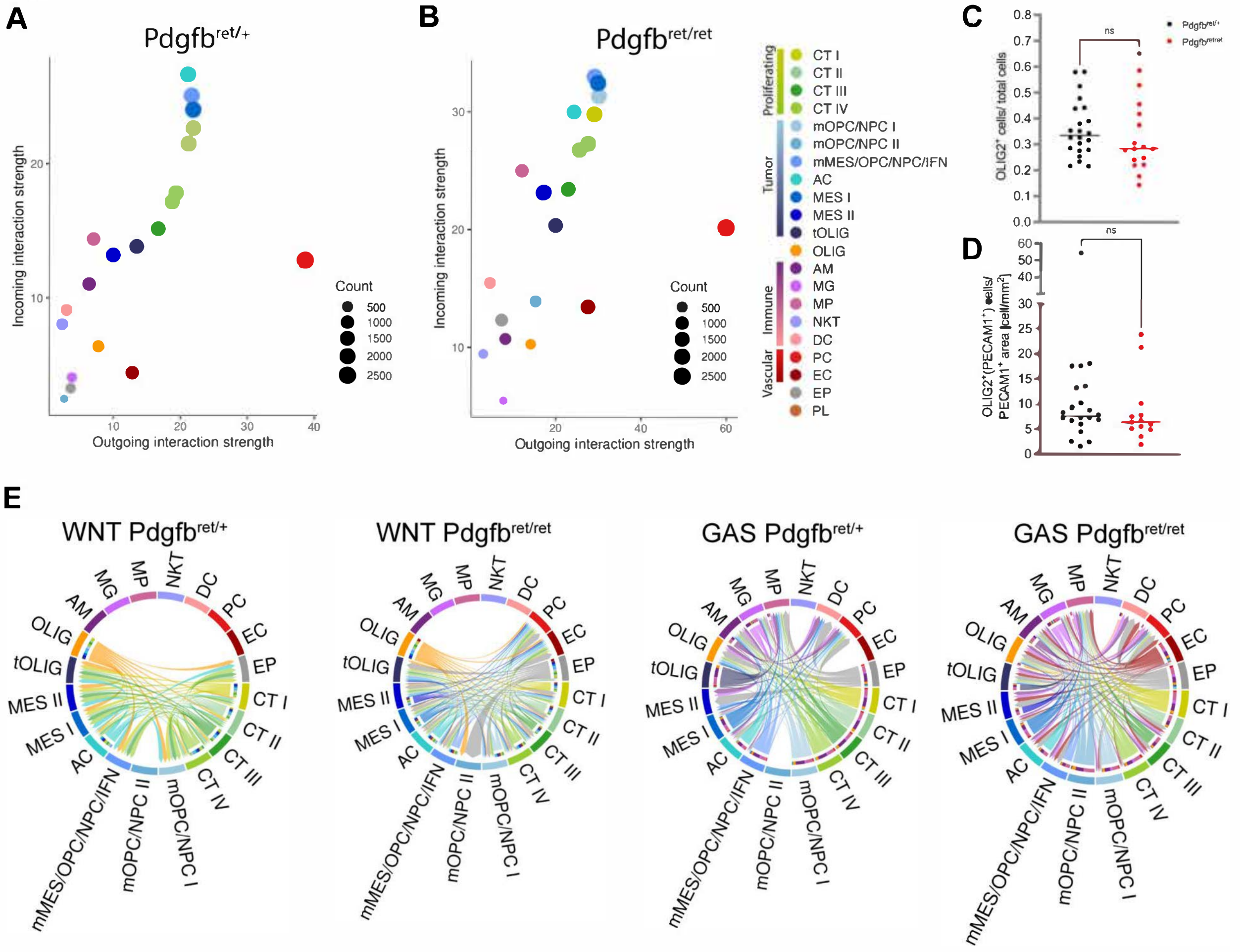

A
Pdgfb ret/+
Incoming interaction strength
Outgoing interaction strength
Count
500
1000
1500
2000
2500
B
Pdgfb ret/ret
Incoming interaction strength
Outgoing interaction strength
Count
500
1000
1500
2000
2500
Proliferating
Tumor
Immune
Vascular
CT I
CT II
CT III
CT IV
mOPC/NPC I
mOPC/NPC II
mMES/OPC/NPC/IFN
AC
MES I
MES II
tOLIG
OLIG
AM
MG
MP
NKT
DC
PC
EC
EP
PL
C
OLIG2+ cells/ total cells
ns
Pdgfb ret/+
Pdgfb ret/ret
D
OLIG2+(PECAM1+) cells/ PECAM1+ area [cell/mm2]
ns
E
WNT Pdgfb ret/+
WNT Pdgfb ret/ret
GAS Pdgfb ret/+
GAS Pdgfb ret/ret

**Figure S4. Description of cell communication patterns and tumor cell vascular co-option parameters in pericyte-deprived glioma, related to Figure 4.**

(A-B) Scatter plots showing the total outgoing and incoming communication probability (incoming or outgoing interaction strength), associated with each cell population for *Pdgfb*$^{ret/+}$ (A) and *Pdgfb*$^{ret/ret}$ (B) glioma derived cells, estimated using CellChat. Dot sizes represent the number of inferred interactions (both incoming and outgoing).

(C-D) Quantification of the abundance of OLIG2$^{+}$ cells (C), and OLIG2$^{+}$ cells in contact with PECAM1$^{+}$ cells, related to the vessel area (D), in the IR.

(E) Chord diagram of selected signaling pathways that show differences between *Pdgfb*$^{ret/ret}$ and *Pdgfb*$^{ret/+}$ tumors, performed with CellChat. WNT pathway (left panels) and GAS pathway (right panels) are differently active in both *Pdgfb*$^{ret/ret}$ and *Pdgfb*$^{ret/+}$ tumors.

ns not significant

### Figure S5 (1)

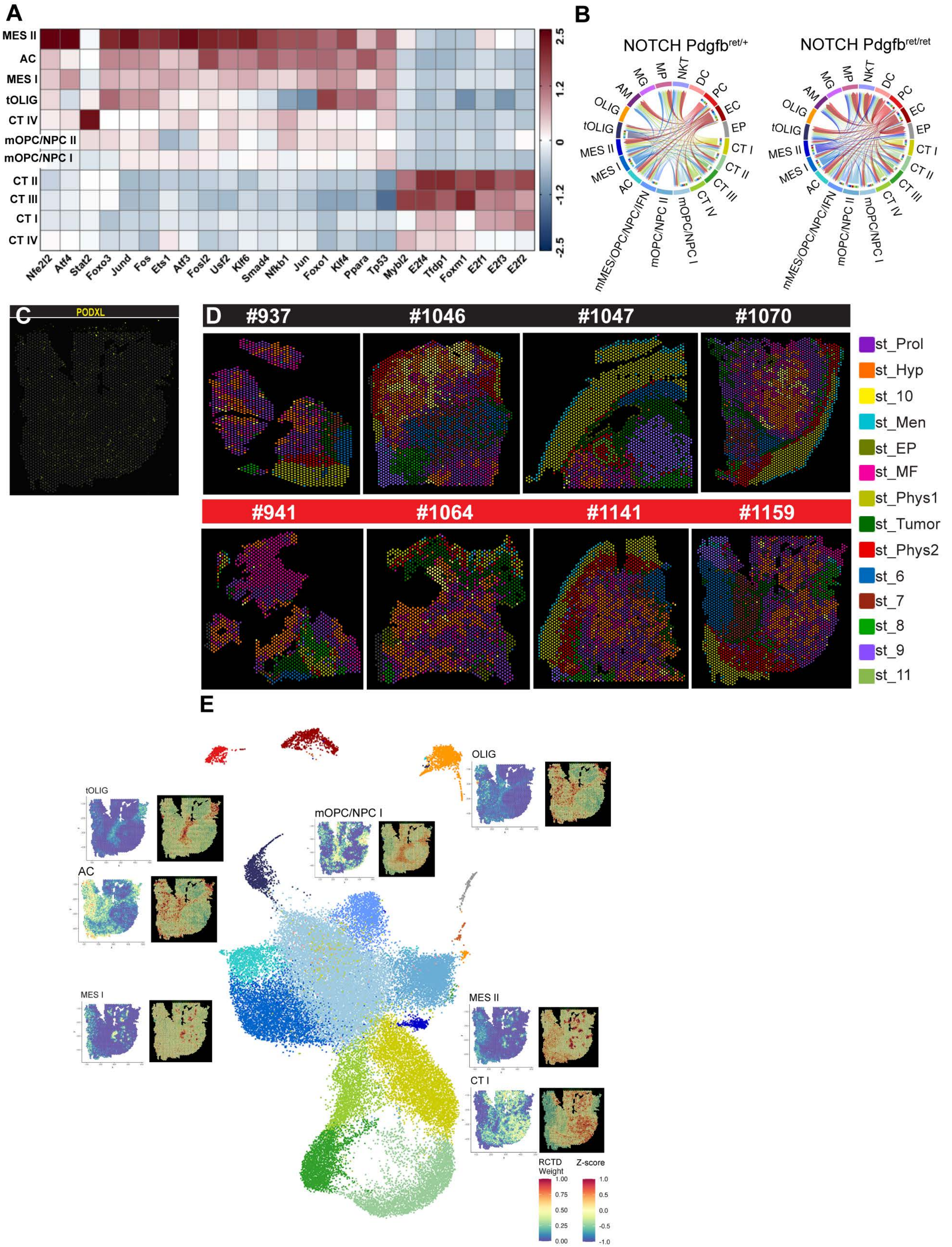

A
MES II
AC
MES I
tOLIG
CT IV
mOPC/NPC II
mOPC/NPC I
CT II
CT III
CT I
CT IV
Nfe2l2
Atf4
Stat2
Foxo3
Jund
Fos
Ets1
Atf3
Fosl2
Usf2
Klf6
Smad4
Nfkb1
Jun
Foxo1
Klf4
Ppara
Tp53
Mybl2
E2f4
Tfdp1
Foxm1
E2f1
E2f3
E2f2
2.5
1.2
0
-1.2
-2.5
B
NOTCH Pdgfb ret/+
NOTCH Pdgfb ret/ret
MG
MP
NKT
DC
AM
PC
OLIG
EC
tOLIG
EP
MES II
CT I
MES I
CT II
AC
CT III
CT IV
mMES/OPC/NPC/IFN
mOPC/NPC II
mOPC/NPC I
C
PODXL
D
#937
#1046
#1047
#1070
#941
#1064
#1141
#1159
st_Prol
st_Hyp
st_10
st_Men
st_EP
st_MF
st_Phys1
st_Tumor
st_Phys2
st_6
st_7
st_8
st_9
st_11
E
OLIG
tOLIG
mOPC/NPC I
AC
MES I
MES II
CT I
RCTD Weight
Z-score
1.00
0.75
0.50
0.25
0.00
1.0
0.5
0.0
-0.5
-1.0

#### Figure S5 (2)

F

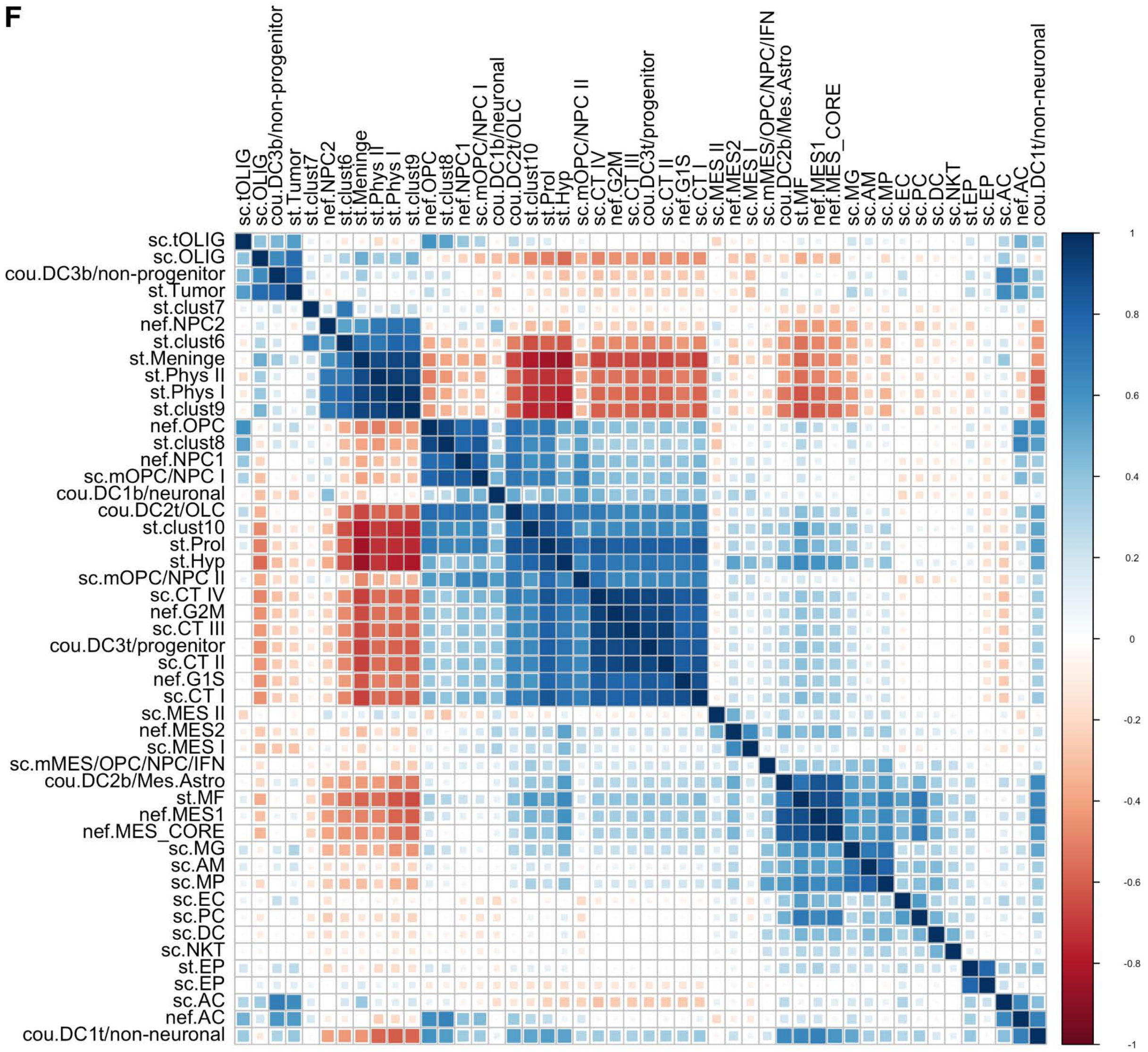


G

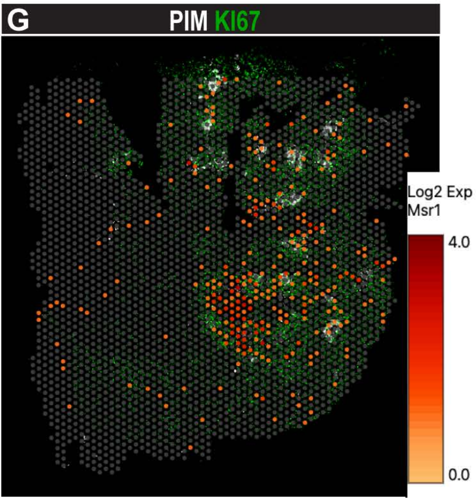

### Figure S5 (3)

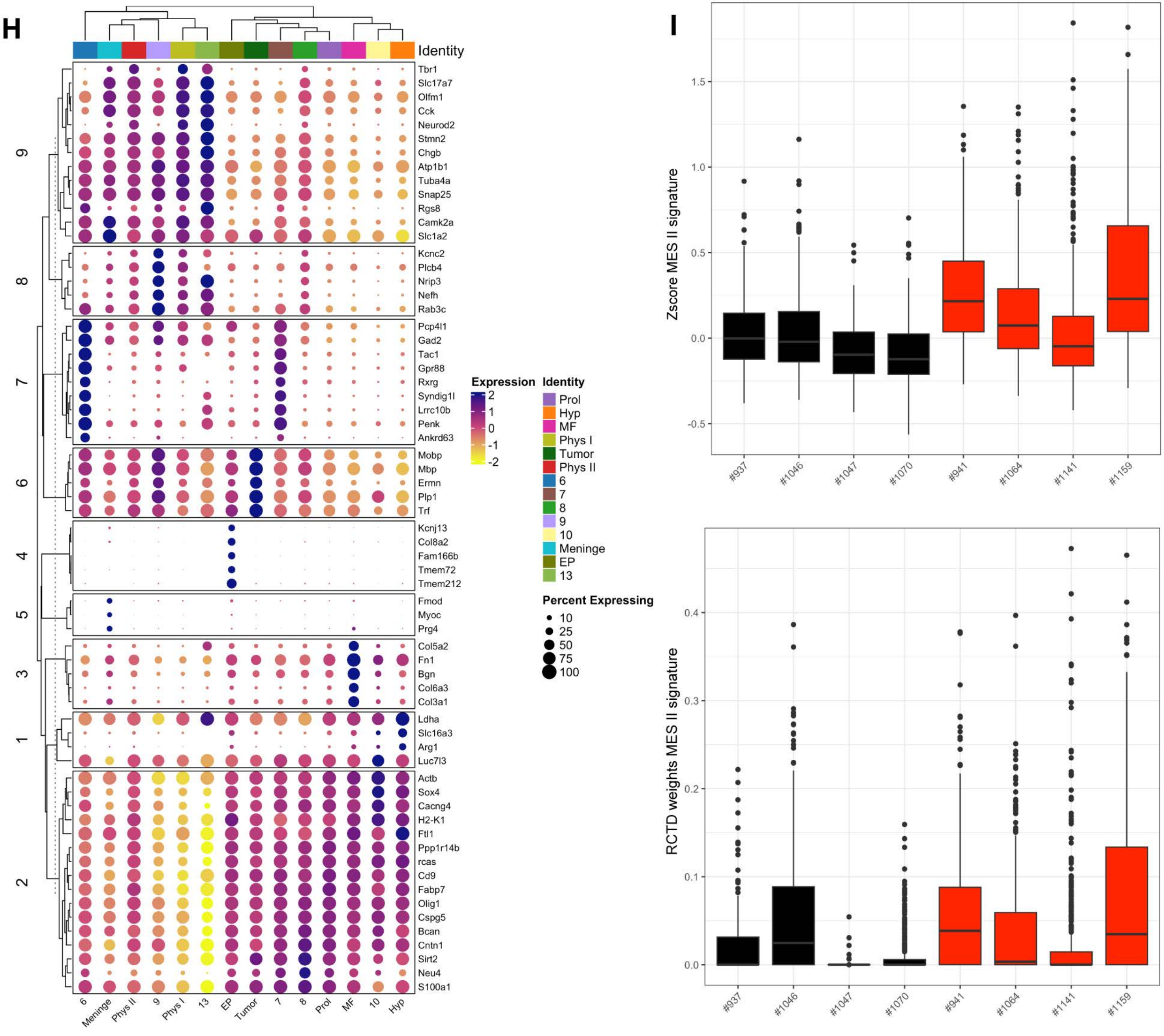
H
Identity
Tbr1
Slc17a7
Olfm1
Cck
Neurod2
Stmn2
Chgb
Atp1b1
Tuba4a
Snap25
Rgs8
Camk2a
Slc1a2
Kcnc2
Plcb4
Nrip3
Nefh
Rab3c
Pcp4l1
Gad2
Tac1
Gpr88
Rxrg
Syndig1l
Lrrc10b
Penk
Ankrd63
Mobp
Mbp
Ermn
Plp1
Trf
Kcnj13
Col8a2
Fam166b
Tmem72
Tmem212
Fmod
Myoc
Prg4
Col5a2
Fn1
Bgn
Col6a3
Col3a1
Ldha
Slc16a3
Arg1
Luc7l3
Actb
Sox4
Cacng4
H2-K1
Ftl1
Ppp1r14b
rcas
Cd9
Fabp7
Olig1
Cspg5
Bcan
Cntn1
Sirt2
Neu4
S100a1
9
8
7
6
4
5
3
1
2
6
Meninge
Phys II
9
Phys I
13
EP
Tumor
7
8
Prol
MF
10
Hyp
Expression
2
1
0
-1
-2
Identity
Prol
Hyp
MF
Phys I
Tumor
Phys II
6
7
8
9
10
Meninge
EP
13
Percent Expressing
10
25
50
75
100
I
Zscore MES II signature
1.5
1.0
0.5
0.0
-0.5
#937
#1046
#1047
#1070
#941
#1064
#1141
#1159
RCTD weights MES II signature
0.4
0.3
0.2
0.1
0.0
#937
#1046
#1047
#1070
#941
#1064
#1141
#1159

**Figure S5. Validation and integration of the scRNA-seq and stRNA-seq data, related to Figure 5.**

(A) Transcription factor activity analysis of the scRNA-seq data using DoRothEA. The heatmap represents the top 25 transcription factor activity scores among the identified cell groups.

(B) Chord diagram of the NOTCH signaling pathway, comparing its activity in *Pdgfb*$^{ret/ret}$ and *Pdgfb*$^{ret/+}$ tumors, performed with CellChat.

(C) VGES capture area of stRNA-seq sample #1159, showing an immunostaining of PODXL (this panel is complementary to Figure 5B).

(D) VGES capture area images of 4 *Pdgfb*$^{ret/+}$ (left panels, black label) and 4 *Pdgfb*$^{ret/ret}$ (right panels, red labels) glioma sections, indicating 14 spatial clusters, identified based on the integration of all 8 samples, using Harmony.

(E) scRNA-seq UMAP plot, together with the VGES of stRNA-seq sample #1159 colored by mean Z-scores (right panels) and normalized RCTD weights (left panels) of selected scRNA-seq cell group signatures.

(F) Clustered correlation matrix showing pairwise Pearson's correlation coefficients of the mean Z-scores from signatures of different cell identity or spatial transcriptomics clusters from the scRNA-seq and stRNA-seq datasets from this study, Neftel et al. (nef) and Couturier et al. (cou).

(G) VGES capture area of stRNA-seq sample #1159, showing the logNormalized expression of *Msr1*, together with immunostaining of KI67 and PIM.

(H) Clustered dot plot showing the scaled average expression of selected top differentially expressed genes across all stRNA-seq clusters.

(I) Boxplots showing the MES II signature Z-scores (upper panel) and inferred proportion of MES II cells by RCTD (lower panel) for the *Pdgfb*$^{ret/+}$ (black bars) and *Pdgfb*$^{ret/ret}$ (red bars) stRNA-seq glioma samples.

### Figure S6

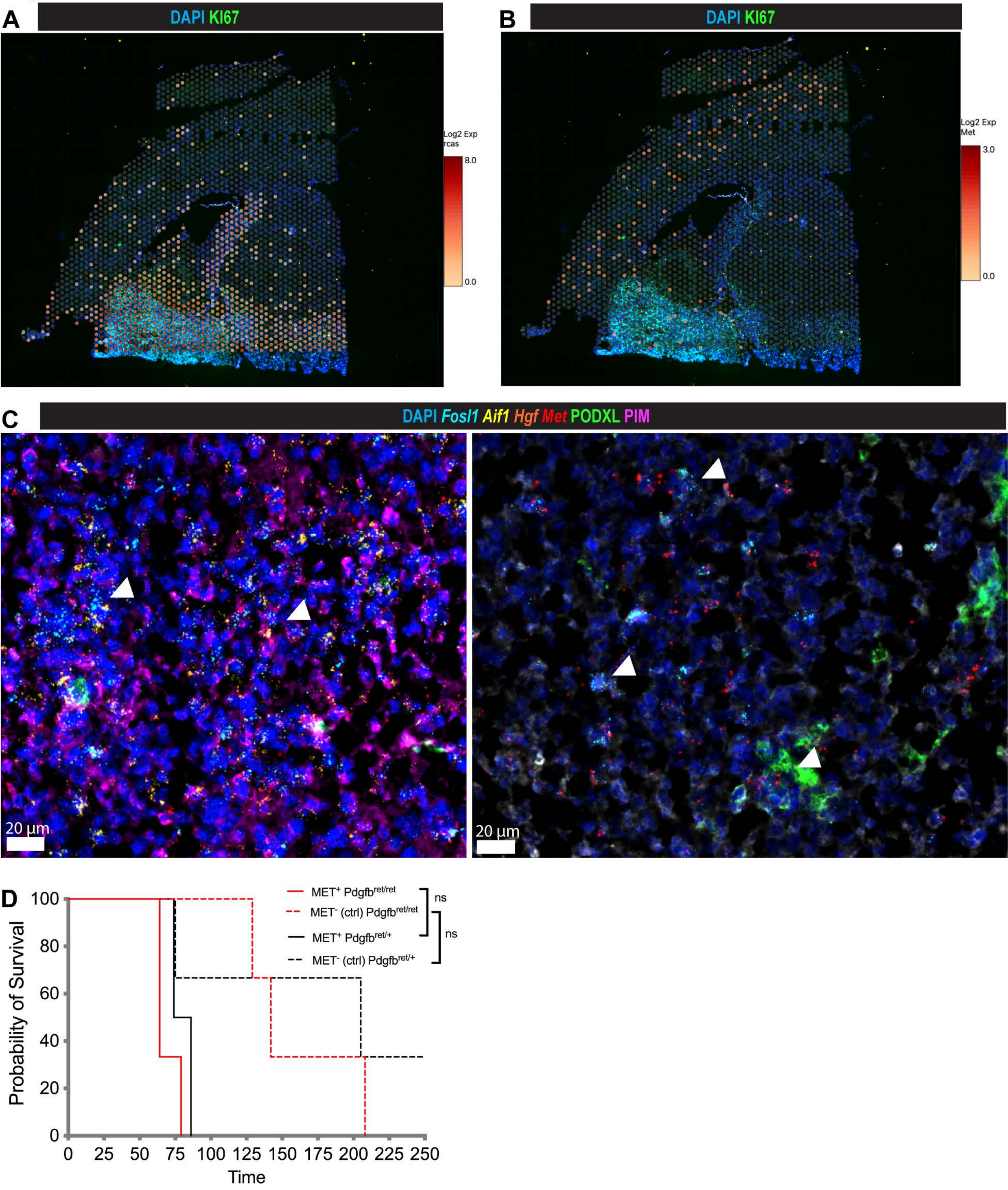

A
DAPI KI67
Log2 Exp
rcas
8.0
0.0
B
DAPI KI67
Log2 Exp
Met
3.0
0.0
C
DAPI Fosl1 Aif1 Hgf Met PODXL PIM
20 µm
20 µm
D
MET+ Pdgfb ret/ret
MET- (ctrl) Pdgfb ret/ret
MET+ Pdgfb ret/+
MET- (ctrl) Pdgfb ret/+
ns
ns
Probability of Survival
100
80
60
40
20
0
0 25 50 75 100 125 150 175 200 225 250
Time

**Figure S6. Localization and *in vivo* assay of MES II cells, related to Figure 7.**

(A-B) VGES capture area of stRNA-seq sample #1047, showing the expression of an RCAS unique sequence (A) and *Met* (B), together with immunostaining of KI67 and PIM.

(C) Combined multiplexed immunostaining-ISH analysis for the detection of *Hgf*, *Aif1*, *Fosl1*, *Met*, PODXL and PIM in PDGFB-induced gliomas, derived from *Pdgfb*$^{ret/ret}$ mice.

(D) Kaplan-Meier curves showing symptom-free survival of mice transplanted with MET$^{+}$ and MET$^{-}$ glioma cells, stratified into *Pdgfb*$^{ret/ret}$ and *Pdgfb*$^{ret/+}$ mouse groups. ns not significant

### Figure S7

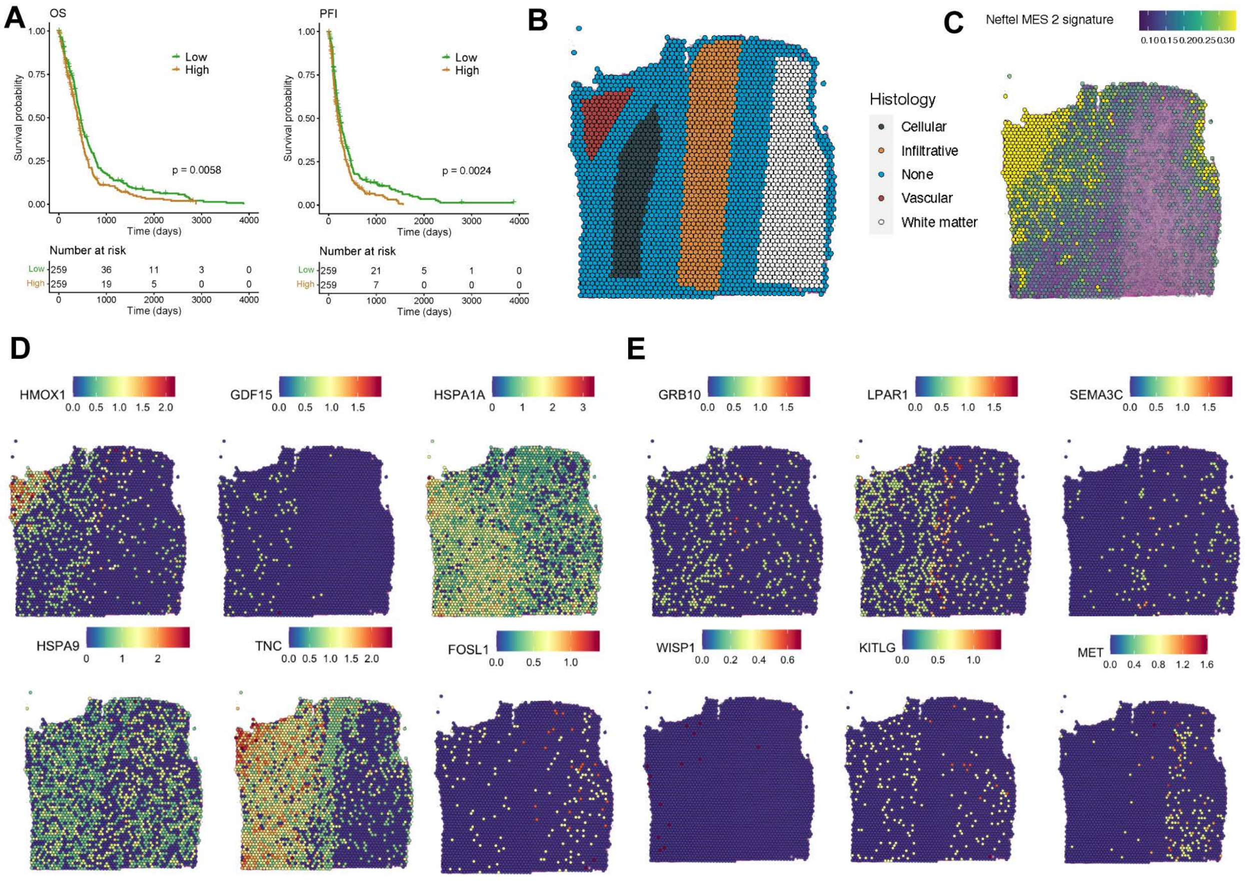

A
OS
PFI
Low
High
Survival probability
Time (days)
p = 0.0058
p = 0.0024
Number at risk
Low 259 36 11 3 0
High 259 19 5 0 0
Low 259 21 5 1 0
High 259 7 0 0 0
B
Histology
Cellular
Infiltrative
None
Vascular
White matter
C
Neftel MES 2 signature
D
HMOX1
GDF15
HSPA1A
HSPA9
TNC
FOSL1
E
GRB10
LPAR1
SEMA3C
WISP1
KITLG
MET

**Figure S7. Spatial characterization of GBM patient samples, based on histopathological and transcriptomics analysis, related to Figure 8.**

(A) The Kaplan-Meier curves show the overall survival (OS) and progression-free interval (PFI) probabilities of the high (brown) and low (green) MES II expression group in the TCGA GBM cohort (n = 518). p-value: log-rank test.
(B) Spatial plot showing the histological morphology assessment in sample #UKF269_T made by Ravi *et al.* (Ravi et al., 2022)
(C) Spatial plot and heatmap showing the MES-like (Neftel et al.) module scoring in sample #UKF269_T.
(D-E) Spatial plots showing the expression of HSS genes (D) and ISS genes (E) in sample #UKF269_T.
